# A dopamine circuit regulates locomotor initiation and persistence in *Drosophila*

**DOI:** 10.1101/2025.11.18.689052

**Authors:** Xinke Wen, Kristin E. Connors, Chris C. Snell, Ryan McGrath, Avirut Mehta, Daisuke Hattori

**Affiliations:** Department of Physiology, UT Southwestern Medical Center; Dallas, TX, 75390, USA; Department of Neuroscience, UT Southwestern Medical Center; Dallas, TX, 75390, USA; Peter O’Donnell Jr. Brain Institute, UT Southwestern Medical Center; Dallas, TX, 75390, USA

## Abstract

Decisions to initiate or terminate locomotion reflect the commitment of an animal to expend energy and thus must be appropriately regulated. Dopaminergic system has been implicated in locomotor regulation but how it controls these decisions remains unclear. Here we show that a dopamine circuit in *Drosophila* mushroom body regulates cue-induced locomotor initiation and termination by integrating locomotor history and current motivation. This circuit consists of the locomotor-initiator mushroom body output neurons, MBON09, and the locomotor-terminator MBON21. Previous locomotor initiation by default suppresses the propensity of future initiation through depression of MBON09 activity by locomotion-sensitive dopamine afferents, preventing redundant action. Locomotor persistence is promoted through combined inhibition of MBON21 by MBON09 and by distinct dopamine afferents that receive fluctuating motivational signals.

Persistent locomotion under high motivational state, in turn, causes dopamine-dependent MBON09 facilitation, reinvigorating locomotor initiation program. Our results revealed a dopaminergic mechanism to transform recent behavior and current motivation into a moment-by-moment internal state that in turn regulates locomotor decisions.

## MAIN

Locomotor ability provides animals with the means to explore the environment for better resources, but with energetic cost. Appropriate regulation of locomotion is therefore primal, especially when hungry animals forage for food with limited energy store. Ethological studies of foraging behaviors across animal species revealed that locomotor regulation during foraging aligns closely with theoretical optima, suggesting that the selective pressure has sculpted neural circuits that function economically to control foraging-related locomotor behaviors ^1–5^.

Dopamine is a key player indicated in locomotor regulation in the central nervous system across species. In both vertebrates and invertebrates, dopaminergic neurons exhibit activities that correlate with locomotion, and manipulation of dopaminergic system exerts various effects on locomotor behaviors ^6–14^. Dopaminergic neurons also play a key role in providing plasticity-inducing teaching signals that underlie associative and reinforcement learning through their responses to reward, punishment or cues that predict them ^15–25^. These different functions of dopamine in controlling learning or locomotion are likely mediated by distinct dopaminergic neuron subclasses and their effects on downstream circuits. While the role in learning has been extensively studied, how dopamine and its circuits control locomotion remains unclear.

We took advantage of the ample resources available for *Drosophila* mushroom body and revealed how different dopamine signals and their downstream circuits affect locomotor decisions. The mushroom body is a central brain structure extensively innervated by dopaminergic neurons (DANs) and has long been studied as a site of olfactory associative learning ^26^. DANs innervate the output lobes of the mushroom body ^27^, in which the Kenyon cells (KCs) encoding odor identities ^28–30^ make excitatory synapses with mushroom body output neurons (MBONs) that bias behaviors ^31–35^. Dendrites of MBONs and axons of DANs divide the lobes into 15 discrete compartments, each containing a unique set of MBON and DAN subclasses ^36,37^. During associative learning, specific DANs innervating specific compartments are activated by reward or punishment (i.e., unconditioned stimulus, or US) and induce plasticity at the active KC-MBON synapses encoding the coinciding odor (i.e., conditioned stimulus, or CS) ^7,38,39^. This compartment-and synapse-specific plasticity results in altered CS response of cognate MBONs, which then drive specific learned behaviors, like avoidance or attraction ^40–44^. Interestingly, as in mammals, some DANs are activated in the absence of US, for example upon locomotion or odor encounter ^7,45,12–14,46^. In this study, we focused on the initiation and termination of odor-induced locomotion in hungry flies—two key locomotor decisions that determine energy expenditure—and revealed a hierarchical mushroom body circuit that coordinately regulates these locomotor decisions by integrating recent locomotor history and current motivation through distinct dopamine-mediated modulatory mechanisms. This concise circuit likely contributes to an energy-efficient foraging behavioral program.

## RESULTS

### Moment-by-moment internal state affects initiation and persistence of odor-induced locomotion

A foraging animal makes a series of locomotor decisions to maximize the potential gain while minimizing the cost. Upon encountering a food-predictive cue like smell or sound of prey, the animal first needs to decide whether to initiate locomotion to explore the environment, and then when to terminate locomotion. These decisions affect the energy spent per cue encounter and thus are likely under coordinated regulation. We devised a simple behavioral assay that examines whether grooming flies initiate locomotion upon exposure to food odors and, if so, how long they persist locomoting. Upon exposure to a food-associated odor, apple cider vinegar (ACV), flies typically stopped grooming and started walking (Fig. 1a). We operationally call this odor-induced locomotion “alerting behavior”, which was quantified using three different metrics—velocity of each fly, frame-by-frame annotation of fly behavior, and variance in pixel intensity across consecutive video frames—that are well correlated with each other (Extended Data Fig. 1b-f). As predicted, food-deprivation enhanced alerting behavior in response to ACV (but not in response to a non-food odor, benzaldehyde (BZA); Extended Data Fig. 2a,b), whereas acute feeding suppressed alerting to ACV (Extended Data Fig. 2c,d), indicating that this behavior represents an initial behavioral response of a foraging fly upon encountering a food-associated cue. Curiously, food-deprived flies did not alert to every presentation of ACV (i.e., trial) (Fig. 1b left). On average, flies alerted in 35% of the ACV trials with slight habituation (43% for the first five ACV trials vs 27% for the next five ACV trials; Fig. 1c). The duration of locomotion upon alerting was also variable, ranging from less than a second to 30 seconds (Fig. 1b,d and Extended Data Fig. 1g). Even with a head-fixed behavioral assay, in which ACV was presented from a set distance and angle consistent across trials, we observed variations both in the likelihood of alerting as well as the duration of ensuing locomotion (alerting in 51% of the trials; locomotion duration ranging from one to 50 seconds; Fig. 1e-g and Extended Data Fig. 1g). Increasing the concentration of ACV by three-fold did not result in more consistent behaviors (Extended Data Fig. 1h). These results suggest that observed variations in alerting behavior are not due to inconsistency in sensory detection and, instead, result from a moment-by-moment internal state of each fly that fluctuates on the sub-minute timescale, providing an experimental system to investigate the underlying regulatory mechanisms.

**Fig. 1:**
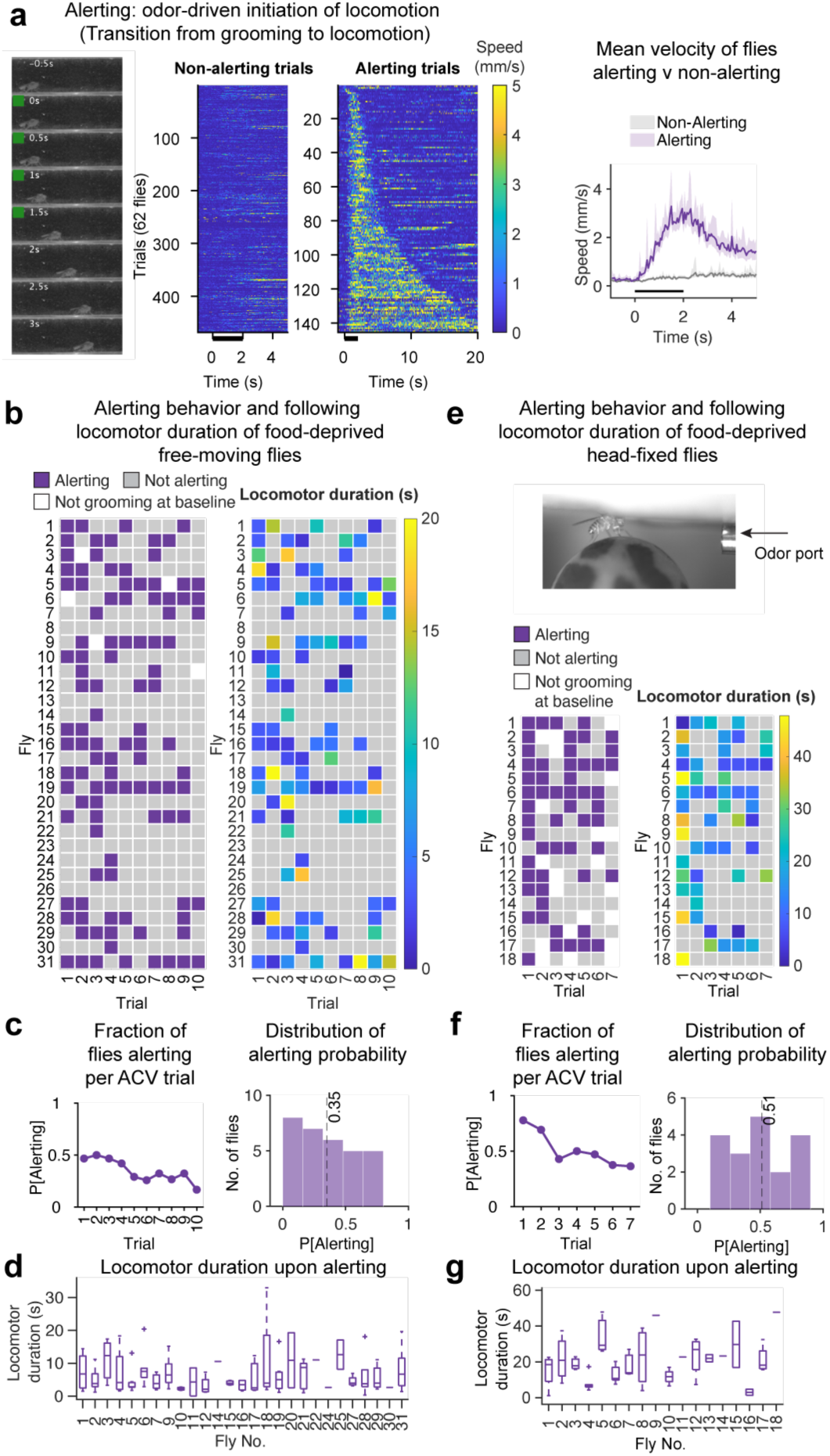
Variable likelihood and duration of alerting behavior in response to apple cider vinegar. (a) Alerting behavior elicited upon apple cider vinegar (ACV) stimulation. Left, example video frames of alerting behavior exhibited by free-moving fly. Green squares indicate presence of odor. Center two panels, speed of flies in alerting vs non-alerting trials. See Methods for definition of alerting. Alerting trials, N=138; non-alerting trials, N=474; from 62 flies. Right, average speed of flies in alerting vs non-alerting trials. Shading indicates 95% confidence interval determined by bootstrapping (see Methods). Black bar indicates odor period (same below). (b) Alerting behavior of food-deprived free-moving flies in response to ACV presentations. Left, trial-by-trial classification of fly behavior; right, duration of continued locomotion upon alerting. (c) Probability of alerting across ACV presentations (left) and across flies (right). Dashed line, average probability of alerting across 31 flies. (d) Distribution of locomotor duration in alerting trials across flies. Flies with no alerting trials are not shown. Boxplot in this and all subsequent panels; box – first and third quartiles, line – median, whisker – data within 1.5x interquartile range, cross – outliers. (e) Alerting behavior of food-deprived head-fixed flies in response to ACV presentations. Top, experimental setup. Bottom, same as (b). (f,g) Data for head-fixed flies as in (c) and (d).

### MBON09 drives locomotor initiation in response to food odor

We first set out to identify circuits that drive alerting behavior in response to ACV. Optogenetic silencing of all KCs, the principal neurons of the mushroom body, by expression of green-light sensitive anion channelrhodopsin GtACR1 ^47,48^ eliminated alerting behavior in response to either ACV or BZA, demonstrating that the mushroom body is necessary for alerting behavior (Fig. 2b-d and Extended Data Fig. 3a-c; MB625C>GtACR1 combining a split-GAL4 driver, MB625C ^49^, with a UAS-GtACR1 transgene; see Extended Data Table 4 for sample size and statistics for this and all subsequent experiments). The alerting behavior is likely elicited through the activation of one or a combination of MBONs that receive KC inputs. Optogenetic activation and silencing screens identified a class of GABAergic MBONs as a key player mediating alerting in response to ACV. These MBONs extend their dendrites to the γ3 and β’1 compartments (Fig. 2e; MB083C split-GAL4 driver labeled MBON09 [MBON-γ3β’1] and MBON08 [MBON-γ3] ^36^; for simplicity, we will describe these neurons as MBON09). Flies expressing a red-shifted channelrhodopsin CsChrimson ^50^ in MBON09 elicited strong alerting behavior in response to red-light stimulation in the absence of odors (Fig. 2f and Extended Data Fig. 4a-c). Conversely, optogenetic silencing of MBON09 using GtACR1 eliminated alerting to ACV, mirroring the behavior observed upon silencing all KCs (Fig. 2g and Extended Data Fig. 3d-f; an additional MBON09 driver—Extended Data Fig. 3g; regardless of whether ACV was delivered to flies that were grooming or standing still—Fig. 2h). Optogenetic silencing of MBON09 also eliminated alerting in response to another food-associated odor from strawberry extract (Fig. 2i). In contrast to KC silencing, however, flies in which MBON09 was silenced still elicited alerting behavior in response to the non-food odor, BZA (Fig. 2g and Extended Data Fig. 3d-f), suggesting that different circuits control alerting in response to food vs non-food odors (see Discussion). These results demonstrate that MBON09 plays an essential role in mediating alerting behavior—a transition from resting to locomotive state—upon encountering food-associated odors.

**Fig. 2:**
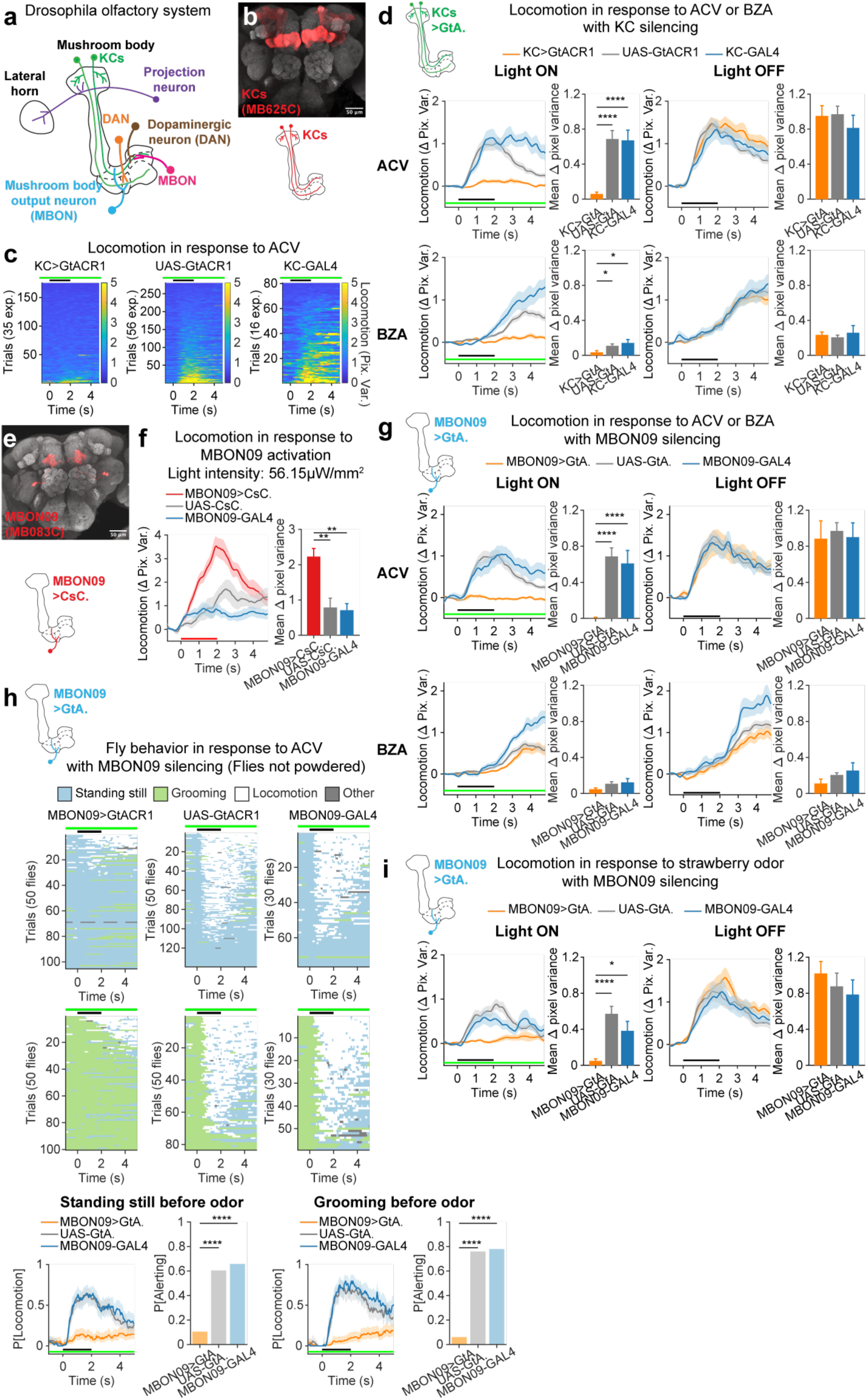
MBON09 mediates the initiation of locomotion in response to food odors. (a) Schematic of fly olfactory circuit. (b) Confocal micrograph of brain showing expression pattern of MB625C split-GAL4 driver (KCs). Neuropil stain by nc82. (c) Effect of optogenetic KC silencing upon locomotion (pixel variance, see Methods) in response to ACV. 2s ACV presentations (black bars) while green light was on continually (green bars). Trials from 35 (KC>GtACR1), 16 (KC-GAL4), 56 (UAS-GtACR1) individual experiments with 4–6 flies per experiment. 5 trials per experiments. (d) Effect of KC silencing upon locomotion in response to ACV or BZA (Δpixel variance, see Methods; see Extended Data Fig. 3a-c for locomotion measured by behavioral annotation). GtA, GtACR1. Mean±SEM in this panel and all subsequent Δpixel variance over time data. Kruskal-Wallis test with post hoc Bonferroni test. *p<0.05; ****p<0.0001. (e) Confocal micrograph of brain showing expression pattern of MB083C split-GAL4 driver (MBON09). (f) Effect of optogenetic MBON09 activation upon locomotion (Δpixel variance). Red bar indicates red light. See Extended Data Fig. 4a-c for experiments with different light intensities. CsC, CsChrimson. Data from 10 (MBON>CsC.), 10 (MBON-GAL4), 10 (UAS-CsC.) individual experiments with 4–6 flies per experiment. Kruskal-Wallis test with post hoc Bonferroni test. **p<0.01. (g) Effect of MBON09 silencing upon locomotion in response to ACV or BZA (Δpixel variance; see Extended Data Fig. 3d-f for locomotion measured by behavioral annotation). Data from 27 (MBON>GtA.), 15 (MBON-GAL4), 56 (UAS-GtA.) individual experiments with 4–6 flies per experiment. Note UAS-GtA. controls are shared with Fig. 3d as experiments conducted on the same days. Kruskal-Wallis test with post hoc Bonferroni test. ****p<0.0001. (h) Effect of MBON09 silencing upon locomotion in response to ACV in flies not powdered. Behavior was manually annotated and trials are sorted by baseline behavior; either standing still or spontaneously grooming. Other behaviors include courtship, proboscis extension, flailing and jumping (see Methods). Line plots, probability of locomotion over time ± 95% confidence interval in this panel and all subsequent probability-over-time data. Probability of alerting is the fraction of trials flies alerted to ACV. Trials from 50 flies (MBON09>GtA.), 30 flies (MBON09-GAL4), 50 flies (UAS-GtA.). Fisher’s exact test. ****p<0.0001. (i) Effect of MBON09 silencing upon locomotion in response to strawberry extract odor. Trials that flies were moving at odor onset are eliminated (see Methods). Data from 19 (MBON09>GtA.), 18 (UAS-GtA.), 19 (MBON09-GAL4) individual experiments with 4–6 flies per experiment. Kruskal-Wallis test with post hoc Bonferroni test. *p=0.027; ****p=1.7E-5.

ACV-induced alerting behavior is modulated on the timescale of hours by feeding states and on the timescale of minutes from trial to trial. The observed variations in alerting behavior may result from variations in MBON09 odor responses. Using calcium imaging with calcium indicator GCaMP8f ^51^, we recorded MBON09 activity while simultaneously monitoring fly behavior (Fig. 3a; see Methods and Extended Data Table 3 for 2-photon imaging parameters). ACV was presented to the fly mounted on an air-supported ball in a resting non-locomotive state, either standing still or grooming. Comparison of average MBON09 responses between starved vs fed flies identified little difference, indicating that the feeding state does not modulate overall MBON09 responses (Extended Data Fig. 5a,b). By contrast, we observed a significant trial-by-trial fluctuation of MBON09 ACV responses in food-deprived flies that correlated with the trial-by-trial variation in alerting behavior (but not merely reflecting locomotion, see below). For example, a strong MBON response in one trial that accompanied alerting was suppressed by 70% in the following non-alerting trial, in which the fly remained still, and then recovered to 87% of the initial response in the next trial, in which the fly alerted again in response to ACV (Fig. 3b and Extended Data Fig. 5c-e). Sorting ACV trials into alerting vs non-alerting trials revealed that calcium responses of MBON09 in alerting trials were on average twice as large as those observed in non-alerting trials (Fig. 3c-e). This association between MBON09 response amplitude and alerting behavior is not due to the slight habituation observed for both MBON09 response and alerting behavior, because strong association remained even after excluding the MBON response and behavior observed in the first ACV trial (Extended Data Fig. 5h). Two additional analyses confirmed the relationship between MBON response and alerting behavior; the alerting probability was significantly higher with stronger MBON odor responses (Fig. 3f) and fly behavior can be accurately classified by MBON09 ACV response amplitudes (Fig. 3g; receiver operating characteristic (ROC) analysis; area under ROC curve (AUC) =0.711, p=0).

**Fig. 3:**
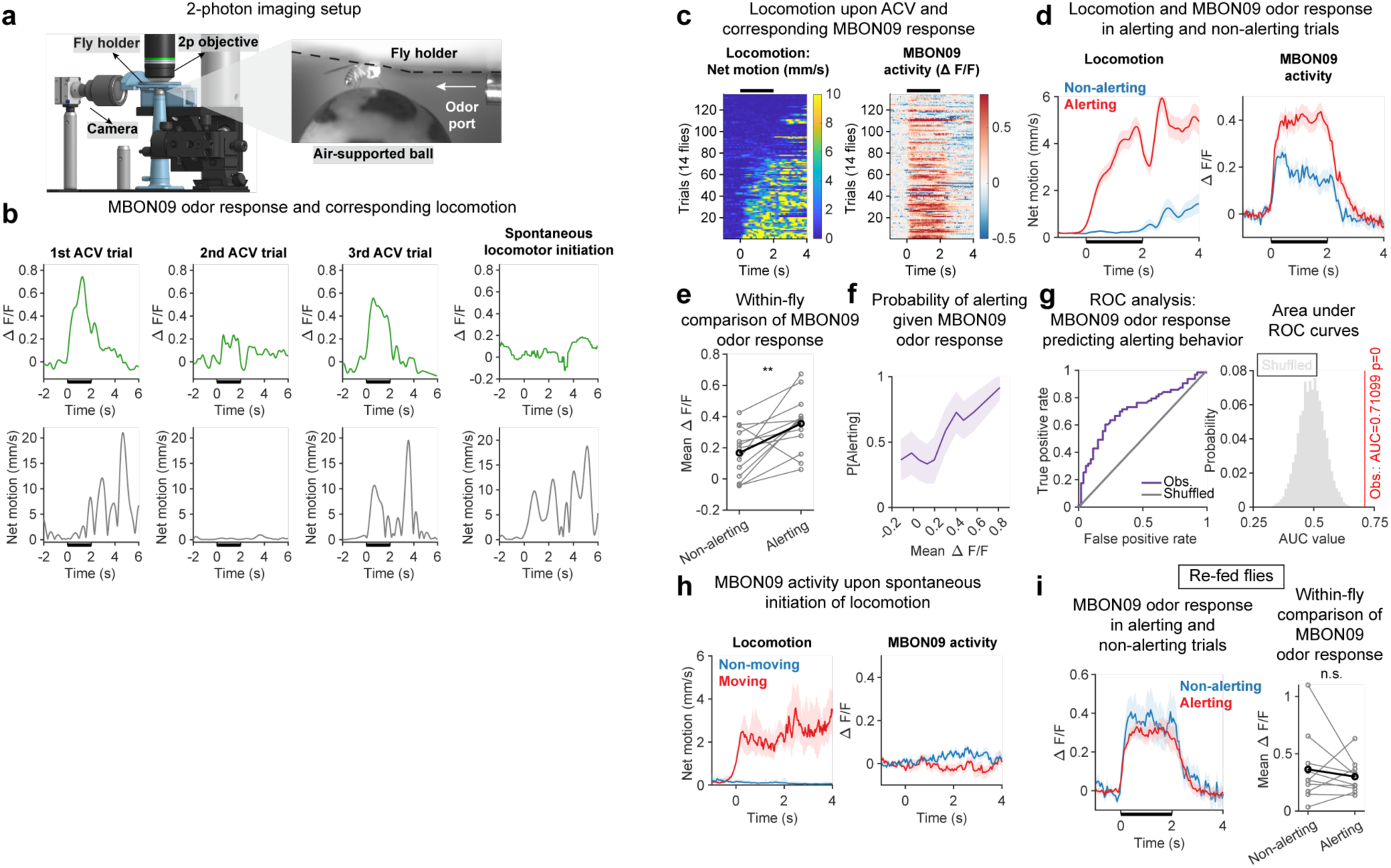
MBON09 odor response predicts the likelihood of initiating locomotion. (a) Experimental setup for 2-photon imaging. (b) Example MBON09 calcium activity (at soma) and corresponding fly locomotion upon three ACV presentations and upon spontaneous initiation of locomotion. ACV trials separated by ∼ 1 min (see Methods). Black bars indicate odor period (same below). (c) Locomotion and corresponding MBON09 activity in response to ACV in food-deprived flies. Trials are sorted based on locomotion. Trials in which fly was moving before odor onset are excluded. (d) MBON09 ACV response in food-deprived flies in alerting vs non-alerting trials (see Methods). Mean±SEM in this panel and all subsequent locomotion and calcium imaging traces. (e) Mean MBON09 ACV responses in alerting vs non-alerting trials; gray lines, individual flies; black line, group average. Flies that did not have both alerting and non-alerting trials are excluded. N=13 flies. Wilcoxon signed-rank test. **p=0.0081. (f) Probability of alerting given MBON09 odor response in food-deprived flies (see Methods). Shading, standard deviation by bootstrapping. (g) Performance of MBON09 response as classifier of alerting behavior for food-deprived flies as evaluated by ROC analysis. Left, ROC curves. Right, permutation test; red line indicates observed area under curve (AUC) value (obs.); gray histogram, distribution of AUC for shuffled data. One-tailed t-test. p=0. (h) MBON09 activity of food-deprived flies upon spontaneous locomotor initiation. N=13 flies. (i) MBON09 ACV response in re-fed flies in alerting vs non-alerting trials, as in (d) and (e). N=10 flies.

Examination of the relationship between MBON response and different behavior parameters (e.g., forward or angular velocity) or odor-delivery angles indicate that strong MBON09 response is observed as long as the fly initiated locomotion in response to ACV without regard to specific parameters of stimulus encounters or resulting locomotor behavior (Extended Data Fig. 6).

The association between MBON09 response and alerting behavior is odor-and feeding-state-specific. First, MBON09 exhibited little response upon spontaneous locomotor initiation (Fig. 3h), indicating that MBON activation requires odor input and that the observed trial-to-trial fluctuation in MBON09 odor response is not a mere consequence of locomotion. Second, MBON09 response upon BZA presentations did not predict alerting to BZA (Extended Data Fig. 5i), consistent with the finding that alerting behavior in response to BZA does not require MBON09. Third, the association between MBON09 ACV response and alerting behavior was eliminated in flies that were acutely fed (Fig. 3i and Extended Data Fig. 5j,k), suggesting that re-feeding shunts the effect of MBON09 upon alerting behavior, likely through modulations that occur at the downstream of MBON09 (see Discussion). Taken together, these results demonstrate that the trial-by-trial fluctuation of ACV-induced MBON09 activity determines the likelihood of whether a food-deprived fly alerts to ACV—an initial decision a hungry fly makes to commit its energy store for foraging exploration. This raises a possibility that suppression and recovery of MBON responses are regulated trial-by-trial to control propensity of alerting behavior.

### Behavioral history modulates MBON09 through dopamine-mediated plasticity to control locomotor initiation

The acute modulation of MBON09 ACV responses likely originates from dopamine-induced changes at KC-MBON synapses, because calcium activity of KCs, a major excitatory input to MBON09, did not differentiate alerting vs non-alerting trials (Extended Data Fig. 5c-g). Previous studies showed that DANs projecting to other mushroom body compartments can affect MBON responses at least in three different ways; direct MBON depolarization ^52^; synaptic depression of active KC-MBON synapses ^7,38,45,42,53,39^; and synaptic facilitation of inactive KC-MBON synapses ^7,38,45,42^. Both MBON09 dendrites and its presynaptic KCs receive input from three classes of DANs; PAM12 (PAM-γ3), PAM13 (PAM-β’1ap), and PAM14 (PAM-β’1m) (Fig. 4a and Extended Data Fig. 7a). We first examined the effect of optogenetically activating these DANs on MBON response. LexA-LexAop binary expression system ^54^ was employed to express GCaMP6f in MBON09 while GAL4-UAS system was used to express CsChrimson in different DANs. These DANs did not directly activate (or inhibit) the MBONs (Fig. 4b) but rather modulated MBON odor responses bidirectionally through plasticity (Fig. 4c,d). ACV-induced MBON activity became strongly suppressed after coincident DAN photoactivation with ACV presentation (93% suppression), whereas MBON response exhibited strong recovery after DAN photoactivation alone without ACV (recovery to 58% of the baseline response). Additional experiments using more specific GAL4 drivers demonstrated that significant modulation of MBON09 ACV response was only observed when PAM12 was manipulated but not when PAM13 and 14 were manipulated (Extended Data Fig. 7b,c). These results demonstrate that PAM12 activity modulates MBON09 odor response bidirectionally based on whether dopamine release coincides with odor.

**Fig. 4:**
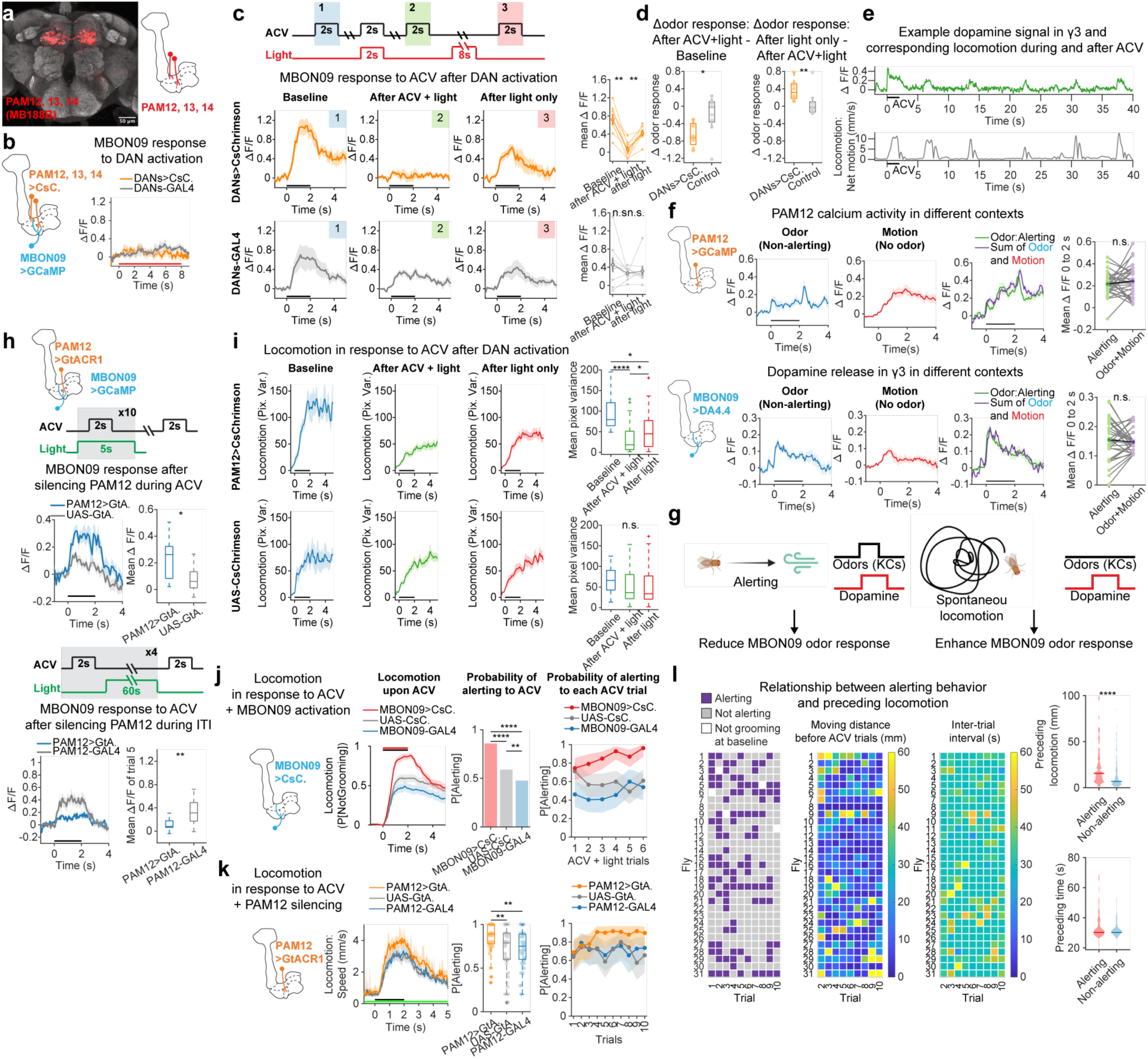
Recent locomotor history biases locomotor initiation through PAM12-MBON09 circuit. (a) Confocal micrograph of brain showing expression pattern of MB188B split-GAL4 driver (PAM12, PAM13 and PAM14). (b) MBON09 calcium activity in response to optogenetic activation of PAM12/13/14. Red bar indicates light period. N=8 (DANs>CsC.) and 9 (DANs-GAL4) flies. (c) MBON09 ACV response before and after optogenetic activation of PAM12/13/14 DANs with or without simultaneous presentation of ACV. Black bars, odor period (same below). Right, mean MBON09 ACV response before and after DAN activation; light orange/gray lines represent individual flies; dark orange/gray lines represent group average; error bars, SEM. N=8 (DANs>CsC.) and 9 (DANs-GAL4) flies. Wilcoxon signed-rank test. **p=0.0078. (d) Changes in MBON09 ACV response upon ACV+DAN activation (left) and upon DAN activation only (right). Wilcoxon rank sum test. *p=0.021; **p=0.0079. (e) Example dopamine signal in γ3 compartment and corresponding locomotion. (f) PAM12 axon calcium activity (top) and dopamine signal in γ3 compartment (bottom) in odor-only (i.e., non-alerting), locomotion-only (i.e., spontaneous locomotor initiation) and odor+locomotion (i.e., alerting) trials. Purple traces in alerting panels and purple dots in right panels represent simple sum of odor-only and locomotion-only traces (see Methods). Only flies with all three types of trials (alerting, non-alerting and spontaneous locomotor initiation) are included. N=12 (GCaMP) and 16 (DA4.4) flies. Wilcoxon signed-rank test. p>0.05. (g) Model of context-dependent modulation of MBON09 ACV response by PAM12. (h) MBON09 ACV response after silencing PAM12 during ACV presentation or during inter-trial intervals (ITI; i.e., silencing during spontaneous locomotion). N=11 (PAM12>GtA.) and 17 (UAS-GtA.) flies for silencing during ACV. N=17 (PAM12>GtA.) and 13 (PAM12-GAL4) flies for silencing during ITI. Wilcoxon rank sum test. *p<0.05; **p<0.01. (i) Head-fixed fly locomotion in response to ACV before and after optogenetic activation of PAM12 DANs. N= 22 (PAM12>CsC.) and 21 (UAS-CsC.) flies. Kruskal-Wallis test with post hoc Bonferroni test. *p<0.05; ****p<0.0001. (j) Effect of MBON09 optogenetic activation upon ACV-induced alerting. Left, locomotor probability over time. Red bar, light; black bar, ACV. Center, probability of alerting across all trials. Right, probability of alerting for each ACV+light trial. Shading in this and all subsequent trial-by-trial probability plots, 95% confidence interval by bootstrapping. Fisher’s exact test. **p<0.01; ****p<0.0001. (k) Effect of PAM12 optogenetic silencing upon ACV-induced alerting. Green bar, light; black bar, ACV. Center, probability of alerting per fly. Right, probability of alerting for each ACV trial. Wilcoxon rank sum test. **p<0.01. (l) Relationship between alerting and preceding locomotion or inter-trial interval. Same dataset as Fig. 1b. Trial-by-trial classification of behavior (left, same as Fig. 1b) and locomotor distance (middle) or inter-trial interval (right) preceding each trial. For the latter two panels, trial 1 is not shown. Right, locomotor distances (top) or inter-trial interval (bottom) preceding alerting vs non-alerting trials. N=92 (alerting) and 183 (non-alerting) trials from 31 flies. Wilcoxon rank sum test. ****p=1.39E-7.

PAM12 can be activated or inhibited in a variety of contexts; previous studies observed its inhibition by sugar reward or mating and its activation by electric shock or self-locomotion ^7,55,13,56^. We therefore examined the activity of PAM12 by calcium imaging of its axon in the context of ACV-induced alerting behavior. We observed PAM12 activity when ACV was delivered as well as when the fly initiated locomotion (Fig. 4e). As some ACV presentations resulted in alerting behavior while others did not, we compared the magnitude of PAM12 calcium responses in three different contexts; when the fly exhibited alerting behavior in response to ACV (odor plus locomotion), when the fly remained non-locomotive upon ACV delivery (non-alerting, or odor only), and when the fly spontaneously initiated locomotion (locomotion only). This analysis revealed that PAM12 activity was modulated additively (Fig. 4f top) and its strongest activation was observed when the fly alerted to ACV. The additive nature of PAM12 activation was also confirmed by imaging dopamine release in the γ3 compartment using a GPCR-activation-based-dopamine sensor (GRAB_DA4.4_) ^57^ (Fig. 4f bottom).

The foregoing results suggest a context-dependent role of PAM12 in the modulation of MBON09 responses (Fig. 4g). Alerting to ACV in one trial accompanies a coincident KC activation and strong dopamine release, which suppresses MBON09 response in the next ACV trial, whereas locomotion in the absence of ACV between trials results in dopamine release without KC activation, thereby recovering MBON09 ACV response. PAM12 silencing experiments confirmed this context-dependent role. Optogenetic silencing of PAM12 (but not PAM13 and 14) during ACV presentation resulted in enhanced MBON responses compared to control flies without GtACR1 expression, which were also observed in the subsequent light-off trials in which PAM12 was no longer silenced, arguing for lasting effect of PAM12 silencing upon MBON responses (Fig. 4h top and Extended Data Fig. 7f). By contrast, silencing PAM12 during the intervals between ACV presentations resulted in decreased MBON responses compared to controls (Fig. 4h bottom and Extended Data Fig. 7g). Thus, our results demonstrate an essential role of odor-and locomotion-sensitive PAM12 in bidirectionally modulating MBON09 ACV responses in a manner reflecting the fly’s recent sensory and behavioral history.

Our observations suggest that PAM12 modulates MBON09 odor response in accord with locomotor history, contributing to the observed variation in alerting behavior. This model posits four predictions. First, forcing strong MBON09 responses to each ACV presentation should generate more consistent alerting behavior. Second, optogenetic activation of PAM12 should bidirectionally modulate alerting behavior in accord with the timing relationship between ACV delivery and photoactivation. Third, eliminating PAM12 activity should result in more consistent alerting behavior. And fourth, locomotor history should predict likelihood of alerting behavior in wild type flies. Consistent with these predictions, we observed that simultaneous photoactivation of MBON09 with ACV presentations resulted in enhanced alerting behaviors (alerting in 86% of trials vs 59% and 48% for controls; Fig. 4j and Extended Data Fig. 8a), that PAM12 photoactivation predictably suppressed or recovered alerting behavior in response to ACV (Fig. 4i and Extended Data Fig. 7d,e), that optogenetic silencing of PAM12 caused more consistent alerting (alerting in 86% of trials vs 70% and 73% for controls; Fig. 4k and Extended Data Fig. 8b,c), and that, in the wild type flies, alerting trials were more likely to be preceded by longer locomotion than non-alerting trials (Fig. 4l). Thus, our results demonstrate that odor-and locomotor-driven PAM12 dopamine activity, through its ability to bidirectionally modulate MBON09 response, affords each fly a mechanism to integrate its own behavioral history into the regulation of locomotor initiation in response to ACV. This mechanism by default diminishes the drive to initiate locomotion in response to the same odor that the fly has previously explored, thereby limiting additional energy cost, unless the fly has locomoted long enough to recover the drive, which likely transfers the fly to a new location. We suggest that the weight of KC-MBON09 synapses, which is updated constantly by dopamine release from PAM12, represents a moment-by-moment internal state that determines the propensity to initiate locomotion.

Regulating the decision to initiate locomotion based on each fly’s recent locomotor history contributes to a more efficient foraging behavioral program (see Discussion).

### MBON09 inhibits the locomotion-terminator, MBON21

Decision to terminate locomotion is the second key step that determines energy expenditure per cue encounter. We next sought to identify circuits that control locomotor persistence upon alerting (e.g., Fig. 1b right). We observed no correlation between MBON09 ACV response (measured either at the soma or the axons) and locomotor duration (Fig. 5e and Extended Data Fig. 10a). However, a behavioral analysis in a small fraction of trials in which MBON09 optogenetic activation was delivered to locomoting flies revealed that MBON09 activation suppressed locomotor termination (i.e., suppressed transition back to grooming; Extended Data Fig. 9a,b). Moreover, prolonged photoactivation of MBON09 resulted in elongated ACV-induced locomotion (Extended Data Fig. 9c,d). These results suggest that MBON09, through its GABA release, directly inhibits neurons that drive locomotor termination, thereby facilitating sustained locomotion while it is activated, for example, during the presence of ACV.

**Fig. 5:**
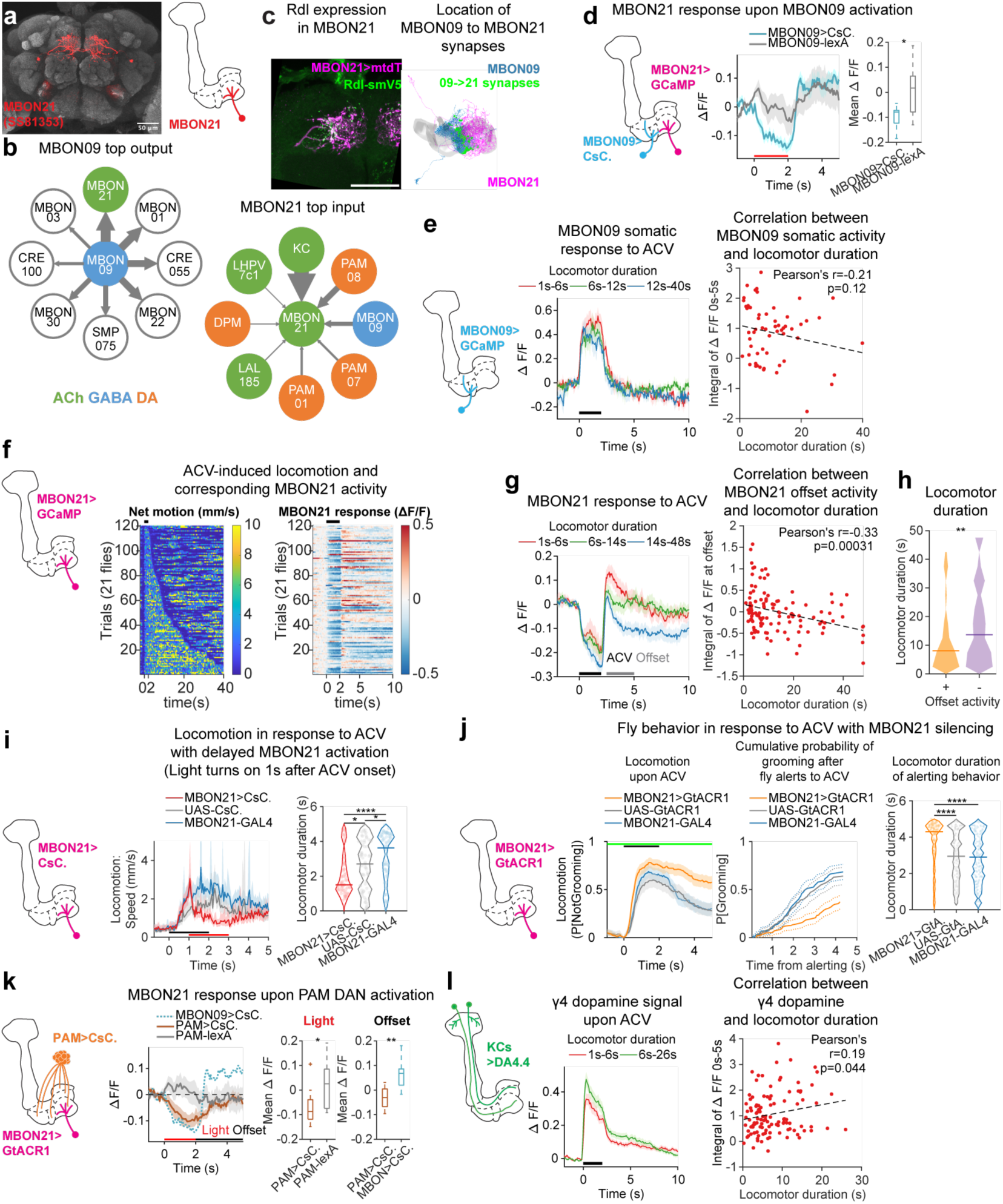
MBON21 integrates inhibitory input from MBON09 and dopamine to regulate locomotor persistence. (a) Confocal micrograph of brain showing expression pattern of SS81353 split-GAL4 driver (MBON21). (b) Downstream neurons of MBON09 (left) and upstream neurons of MBON21 (right). Size of arrows sale with number of synapses. See Extended Data Table 1 and 2 for connection#. (c) Left, expression pattern of GABA_A_ receptor, Rdl, in MBON21 (green, see Methods). MBON21 is labeled with myristoylated tdTomato (magenta, mtdT). Right, MBON09-to-MBON21 synapses (green), MBON09 (blue), and MBON21 (magenta) from a connectome dataset. Gray mesh, mushroom body γ lobe. Scale bar, 50 microns. (d) Effect of MBON09 optogenetic activation upon MBON21 calcium activity. Red bar, light (same below). N=12 (MBON09>CsC.) and 9 (MBON09-lexA) flies. Wilcoxon rank sum test. *p=0.036. (e) Relationship between MBON09 ACV response (soma) and resulting locomotor duration in alerting trials (see Methods). Black bar, ACV (same below). N=19 (1-6 s), 19 (6-12 s), 19 (12-40 s) trials from 14 flies. (f) Locomotion and corresponding MBON21 response to ACV presentations in alerting trials. Trials are sorted by locomotor duration upon alerting. N=21 flies. (g) Relationship between MBON21 ACV response (dendrite) and resulting locomotor duration in alerting trials. N=38 (1-6 s), 39 (6-14 s), 39 (14-48 s) trials from 21 flies. MBON21 offset activity used for right panel (gray bar, 2.5-5s from ACV onset). (h) Sign of MBON21 offset activity and locomotor duration in alerting trials. N=47 (+) and 69 (-) trials. Solid lines indicate group median (same for Fig. 5j). Wilcoxon rank sum test. **p=0.004. (i) Effect of delayed MBON21 optogenetic activation upon locomotion in response to ACV. Dots in violin plots represent trials (same for all violin plots). N=22, 65 (flies, trials; MBON21>CsC.); 17, 72 (MBON21-GAL4); 20, 65 (UAS-CsC.). Wilcoxon rank sum test. *p<0.05; ****p<0.0001. See Extended Fig. 9f for effect of MBON21 activation on alerting probability. (j) Effect of MBON21 silencing upon ACV-induced alerting. Left, locomotion probability (annotation). Center, cumulative probability of resuming grooming after alerting. Solid line indicates mean; dashed line indicates lower and upper confidence bounds for survival function (see Methods). Right, locomotor duration in alerting trials (see Methods). N=26, 216 (flies, trials; MBON21>GtA.); 19, 183 (MBON21-GAL4); 20, 172 (UAS-GtA.). Kruskal-Wallis test with post hoc Bonferroni test. ****p<0.0001. See Extended Fig. 9e,h for effect of MBON21 silencing on locomotor initiation. (k) Effect of PAM DAN optogenetic activation upon MBON21 calcium activity. CsChrimson was expressed by R58E02-lexA driver that labels a subset of PAM DANs, mainly those innervating γ4 and γ5 compartments. Dotted blue line in trace plot is mean MBON21 response upon MBON09 activation (data from Fig. 5d). N=10 (PAM>CsC.), 10 (PAM-lexA) and 12 (MBON09>CsC.) flies. Wilcoxon rank sum test. *p=0.026. **p=0.0011. (l) Relationship between γ4 dopamine signal in response to ACV and resulting locomotor duration in alerting trials. N=56 (1-6 s) and 55 (6-26 s) trials from 36 flies.

We found that MBON21 (also known as MBON-γ4γ5, whose dendrites extend into the γ4 and γ5 mushroom body compartments; Fig. 5a), the top downstream target of MBON09, plays a role in termination of ACV-induced locomotion. MBON09 provides the third most numerous input to MBON21 by type, and it is the only GABAergic neurons among the top 30 MBON21 inputs (Fig. 5b and Extended Data Table 1-2; Data from the hemibrain connectome ^58,59^). We first confirmed the likely GABAergic and inhibitory connections between MBON09 and MBON21 using both anatomical ^60^ and functional imaging experiments. A canonical fly GABA_A_ receptor, Rdl, localized in MBON21 at the boundary between γ3 and γ4 compartments, closely matching the location of MBON09-MBON21 synapses in the EM images (Fig. 5c) and optogenetic activation of MBON09 inhibited MBON21 (Fig. 5d). We then tested the role of MBON21 during ACV-induced locomotion. Acute optogenetic activation of MBON21 in the middle of odor delivery blunted ACV-induced locomotion (Fig. 5i and Extended Data Fig. 9g), whereas photoinhibition of MBON21 resulted in longer ACV-induced locomotion (Fig. 5j and Extended Data Fig. 9h). These results identified MBON21 as a crucial player in controlling locomotor persistence upon ACV-induced alerting and revealed the existence of locomotor-MBON hierarchy comprising direct inhibition of locomotion-terminator MBON21 by locomotion-initiator MBON09.

### Motivational dopamine signal modulates MBON21 inhibition and locomotor persistence

Examination of MBON21 calcium activity in wild type flies revealed that MBON21 was inhibited upon ACV presentations (Fig. 5f), likely resulting from the inhibitory input from MBON09. Curiously, in contrast to MBON09 activation, which was observed only during ACV delivery period, MBON21 inhibition was sustained past ACV delivery period in some trials following a brief rebound activation at ACV offset (Fig. 5f,g). This sustained MBON21 inhibition scaled with locomotor duration; locomotor duration was significantly longer when MBON21 was inhibited after the odor offset (Fig. 5h). An analysis of the relationship between MBON21 offset activity and the duration of locomotion revealed a negative correlation (Fig. 5g right, Pearson’s correlation coefficient r=-0.33), and an ROC analysis indicated that the locomotor duration distinguishes whether the offset activity of MBON21 was above or below the baseline activity observed before ACV delivery (Extended Data Fig. 10b; AUC=0.64, p=0.0023). We noted that while MBON21 inhibition scaled with locomotor duration, it did not determine the precise duration of locomotion (Extended Data Fig. 10c), indicating that MBON21 inhibition is permissive for persistent locomotion. Combined with the behavioral observation (Fig. 5i,j), these results demonstrate that the inhibition of MBON21 during and after ACV delivery promotes the persistence of locomotion.

The observation that MBON21 inhibition persists past odor period indicates the presence of an additional inhibitory input to MBON21 other than MBON09 that contributes to facilitating locomotor persistence. An attractive candidate is the cognate DANs providing extensive inputs into MBON21 (PAM01, PAM07 and PAM08, also known as PAM-γ5, PAM-γ4<γ1γ2 and PAM-γ4; Fig. 5b, Extended Data Fig. 10d and Extended Data Table 2). Previous studies showed that these DANs are activated in numerous motivating contexts, such as upon delivery of appetitive odors, sugar, or water reward and upon goal-directed movement ^61,62,7,13^. Using the GRAB dopamine sensor, we recorded the DAN activity in the γ4 compartment (PAM07/08) during ACV-induced alerting. We observed a trial-by-trial variation in the ACV-induced dopamine release that was positively correlated with locomotor duration (Fig. 5l, Extended Data Fig. 10e,f), suggesting a role of these PAM DANs in regulating locomotor persistence. Consistent with this idea, photoactivation of PAM DANs resulted in sustained inhibition of MBON21, distinct from MBON09-mediated inhibition accompanied by immediate rebound excitation (Fig. 5k). These results indicate that DAN activity facilitates sustained MBON21 inhibition, thereby prolonging locomotor persistence.

The correlation between the odor-evoked dopamine release in the γ4 compartment and ensuing locomotor persistence suggests that PAM07/08 may acutely modulate their ACV response to reflect current locomotor motivation. A previous study showed that the odor response of these DANs are enhanced by food-deprivation, further supporting the idea of modulation by current motivational state ^13^. CRE011, a neuron upstream of PAM07/08, is well positioned to modulate PAM odor response in accord with locomotor motivation, as it integrates inputs from a premotor neuropil, lateral accessary lobe (LAL), and from multiple mushroom body compartments through different MBONs (Fig. 6a, Extented Data Fig. 10g) ^49^. Calcium imaging of CRE011 revealed strong activation induced by ACV, the amplitude of which predicted locomotor persistence (Fig. 6c, d). Interestingly, we also observed a significant fluctuation of CRE011 baseline activity, which did not correlate with observable behaviors (Fig. 6b).

**Fig. 6:**
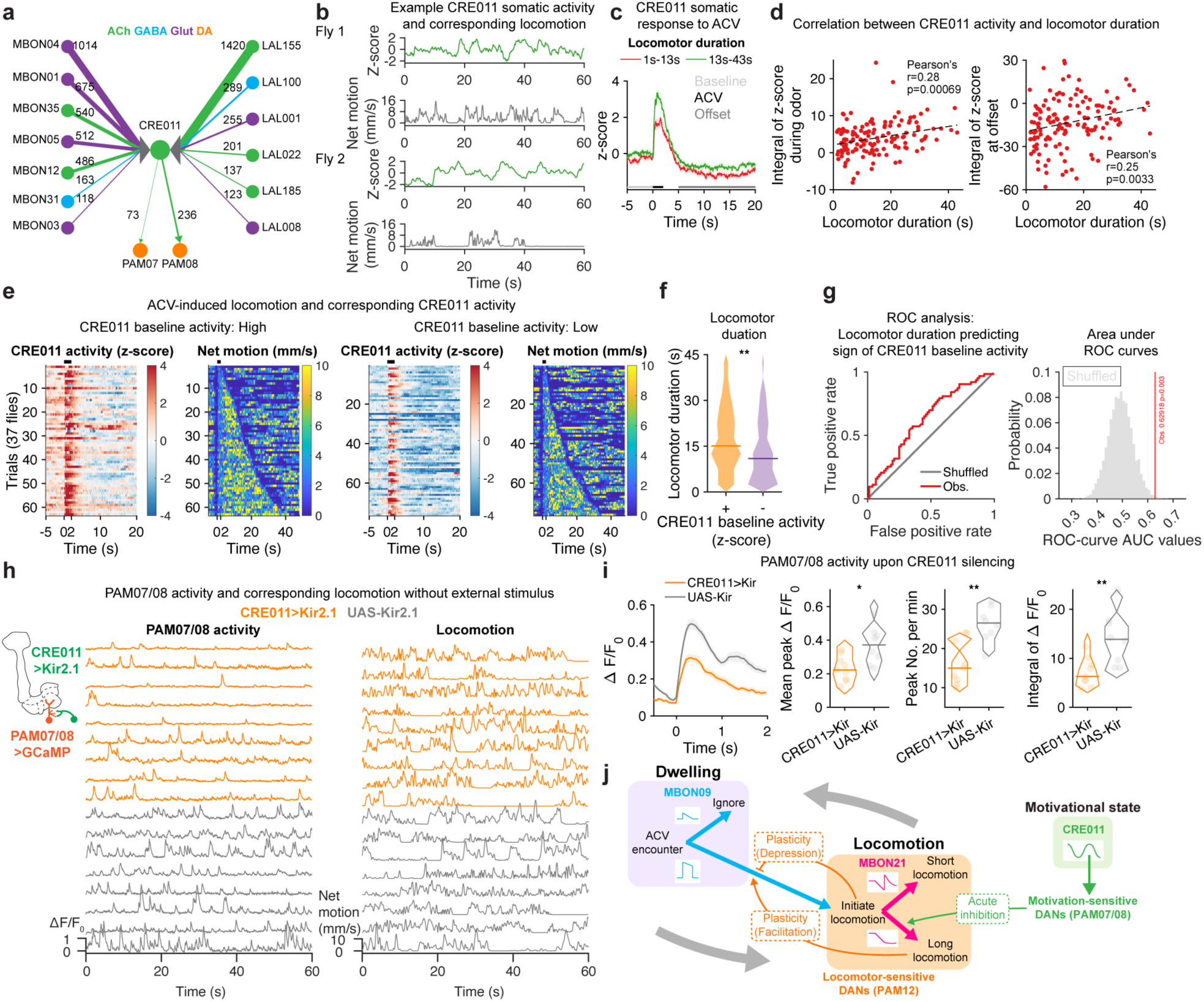
CRE011 conveys motivation signals to PAM07/08. (a) Input connections to CRE011and CRE011 output connections to PAM07/08. Individual MBONs and LALs with more than 100 inputs to CRE011 are listed. (b) Example CRE011 somatic activity and corresponding locomotion. (c) Relationship between CRE011 activity and resulting locomotor duration in alerting trials. CRE011 activity is z-scored using baseline activity before any odor presentations. N=70 (1-13 s), 69 (13-43 s) trials from 37 flies. (d) Correlation between CRE011 activity and locomotor duration in alerting trials, taking integral of neural activity over odor period (left) and offset period (right; 5–20s from odor onset). (e) Locomotion and corresponding CRE011 response to ACV presentations in alerting trials. Trials are separated based on CRE011 baseline activity (5s before odor) and sorted by locomotor duration upon alerting. N=37 flies. (f) Sign of CRE011 baseline activity and locomotor duration in alerting trials. N=63 (+) and 76 (-) trials. Wilcoxon rank sum test. **p=0.0089. (g) Performance of locomotor duration as classifier of the sign of CRE011 baseline (5s before odor) activity as evaluated by ROC analysis. Left, ROC curves. Right, permutation test; red line indicates observed area under curve (AUC) value (obs.); gray histogram, distribution of AUC for shuffled data. One-tailed t-test. p=0.003. (h) Effect of CRE011 silencing upon PAM07/08 calcium activity (left) and locomotion (right) (orange, SS45245>Kir2.1, R58E02>GCaMP6f; gray, EmptyB>Kir2.1, R58E02>GCaMP6f). N=9 (CRE011>Kir) and 8 (UAS-Kir) flies. (i) Comparison of PAM07/08 calcium transient amplitude and frequency in CRE011 silenced flies and control. Dots in violin plots indicate flies. Wilcoxon rank sum test. *p<0.05; **p<0.01. (j) Role of identified circuit in locomotor decisions. Activation of MBON09 controls locomotor initiation whereas inhibition of MBON21 contributes to locomotor persistence. PAM12 reflects locomotor history upon MBON09 responses through bidirectional synaptic plasticity whereas PAM07/08 facilitates locomotor persistence through MBON21 inhibition. CRE011 activity state represents moment-by-moment motivation and affects locomotor duration through PAM07/08.

Stimulation with ACV when CRE011 activity was in a high state resulted in longer locomotion than when it was in a low state (Fig. 6e-g). This effect is likely supported by the influence of CRE011 upon PAM07/08 excitability, as silencing CRE011 significantly reduced both frequency and amplitude of spontaneous calcium transients observed in PAM07/08 (Fig. 6h,i). Thus, our results suggest that the fluctuation in CRE011 activity, which may originate from the LAL circuit, conveys a moment-by-moment motivational state of the fly and affects ensuing locomotor persistence through the PAM07/08-MBON21 circuit, providing a mechanism to reflect current motivation into locomotor persistence.

## DISCUSSION

In this study, we identified a dopamine-modulated hierarchical circuit that regulates locomotor initiation and persistence in response to food odors (Fig. 6j). This circuit consists of two classes of MBONs, the locomotor-initiator MBON09 and the locomotor-terminator MBON21, each tuned by a different dopamine mechanism. Dopamine signals reflect behavioral history or current motivation upon MBON activity, thereby contributing to determining a moment-by-moment internal state of the fly that affects locomotor decisions. Together, our study has established a role of mushroom body and its dopaminergic afferents in controlling locomotor decisions of foraging flies. Distinct dopamine modulations afford this circuit the ability to integrate recent locomotion and current motivation, which likely contributes to more economical foraging.

### Specificity of MBON09 function for food odors

Our results revealed an apparent odor specificity in generating alerting behavior through MBON09. Alerting behavior induced by ACV requires MBON09, whereas alerting in response to BZA does not (Fig. 2g). Moreover, ACV-induced MBON09 response predicts alerting behavior, whereas BZA-induced MBON09 response does not predict alerting (Fig. 3d,e, Extended Data Fig. 5i). These results suggest that MBON09 activity is neither necessary nor sufficient to induce alerting in response to BZA. How could this apparent odor specificity arise and how could it contribute to odor-driven locomotion? First, alerting behavior in response to ACV vs BZA must be mediated by different neural pathways. Indeed, previous studies showed that alerting to BZA is mediated by the MBONs of the α’3 mushroom body compartment ^45^. This division of labor results in activation of different downstream neurons that facilitates distinct patterns of locomotion in response to different classes of odors, like upwind vs downwind locomotion observed in response to attractive (e.g., ACV) vs aversive (e.g., BZA) odors ^63,64^. The odor-compartment specificity also allows alerting behavior induced by different odors to be modulated by distinct population of DANs encoding different external and internal states. This circuit layout aligns the ethological significance of each odor with specific modulatory signals that convey relevant states. Second, the observation that optogenetic activation of MBON09 is sufficient to cause alerting (Fig. 2f), whereas BZA-induced MBON09 activation is not (Extended Data Fig. 5i), indicates that BZA must close the “gate” through which the effect of MBON09 activation propagates to downstream alerting circuit. Thus, there is likely an additional odor-specific neural element sensitive to BZA that converges at the downstream of MBON09.

Candidates include neurons in another olfactory center, lateral horn, which maintains the odor valence information ^65–69^, as well as other MBONs that target common downstream neurons with MBON09. It is also noteworthy that acute feeding results in dissociation of MBON09 response amplitude and alerting behavior (Fig. 3i, Extended Data Fig. 5j,k). This observation indicates that feeding state, in addition to sensory cues, can gate the outcome of MBON09 activation on mediating alerting behavior. Future studies will reveal how the sensory information and the feeding state are integrated in MBON09 downstream circuits.

### Dopamine – reflecting history and current motivation on locomotor decisions

DANs have long been studied in the context of associative learning, in which their response to US and the resulting plasticity of KC-MBON synapses are thought to underlie learned behaviors. For example, DANs in the γ1 compartment respond to aversive US, and paired activation of γ1 DANs with an odor CS induces long-term depression of MBON CS responses, which contributes to learned avoidance ^38^. By contrast, when activated alone, DANs can erase learned avoidance ^70,71,38^. Our results indicate that odor-and locomotor-sensitive dopamine activity in PAM12 regulates KC-MBON09 synapses in a similar manner to that observed in the context of associative learning. Strong PAM12 activation coinciding KC activation during ACV-induced alerting behavior depresses MBON09 ACV response, resulting in decreased likelihood of alerting upon next ACV encounter. By contrast, PAM12 activity upon locomotion in the absence of odor facilitates MBON09 ACV response, resulting in recovered alerting drive. Thus, the PAM12-MBON09 circuit forms a short-term association between the sensory experience and its resulting locomotor response, which is dissociated by locomotion in the absence of sensory stimuli. In this context, flies’ locomotor behavior serves as a US through PAM12 activity, and the same synaptic plasticity rule employed for associative learning is co-opted to reflect history of self-generated locomotion upon future locomotor behavior. By contrast, the effect of PAM07/08 DANs on MBON21 activity is acute and inhibitory, reflecting the current locomotor motivation upon locomotor persistence. Possible mechanism for this PAM-mediated inhibition includes direct inhibitory effect of dopamine and inhibition by co-released GABA ^72,73^. Current locomotor motivation that affects PAM07/08 activity is at least in part conveyed by the activity state of CRE011, which fluctuates on sub-minute timescale. An attractive possibility is that CRE011 activity state is determined by integrating inputs from other MBONs that reflect past experiences and inputs from premotor circuits in the LAL, whose activity encodes premotor state that may correspond to latent locomotor motivation. Finally, the acute effect of dopamine on locomotor persistence identified in our studies is reminiscent to the acute effects of mammalian dopaminergic activation upon locomotion ^9,10^. It will be interesting to examine whether mammalian dopamine system is also employed in reflecting animal’s past self-locomotion upon dictating future locomotor behaviors through dopamine-induced plasticity, as observed for PAM12-MBON09 circuit.

### Contributions of MBON09/MBON21 circuit on foraging behavior

Food-deprivation specifically enhances alerting to ACV, suggesting that alerting to ACV may comprise an initial behavioral response upon food odor encounters in foraging flies. ACV typically induces upwind locomotion ^64^. However, even under condition in which biased upwind turning was induced, we did not observe correlation between MBON09 odor response and the directionality of locomotion (Extended Data Fig. 6). Instead, our findings suggest that MBON09 only acts as an initiator of locomotor cascade when the fly encounters food odors in an awake resting state. Specific movement parameters may be controlled in parallel or downstream circuits, such as those that mediate wind-directed navigation ^74,75^. In this context, it is noteworthy that MBON09 as well as MBON21 provide significant inputs into fan-shaped body tangential neurons that likely affect navigation behaviors ^37,76,35^. In this manner, MBON09 activation may contribute to inducing a general foraging locomotor state in response to ACV, while other circuits control specific behavioral strategies to locate the food source ^77^. It is also noteworthy that previous studies focusing on flies in different initial behavioral states, like running or sleeping, identified roles of different mushroom body compartments in mediating ACV-induced behavior ^78,79^. Thus, multiple mushroom body compartments may be recruited to coordinately execute a robust foraging program in response to food odors.

The regulation of locomotion by MBON09/MBON21 circuit likely contributes to an efficient foraging strategy. First, our observation that strong inhibitory input from MBON09 to MBON21 is sufficient to prevent locomotor termination (Extended Data Fig. 9a-d) suggests that this hierarchical structure facilitates continued locomotion under a constant presence of ACV, which likely signals the proximity of food source. This hierarchy may therefore contribute to limiting locomotion preferentially to when a food source is nearby. Second, diminished alerting propensity upon previous alerting may circumvent an additional energy cost by acutely devaluing redundant action in response to the same sensory information that the fly has recently explored. The same action becomes re-valued upon sustained locomotion, for example, in a high-motivation state that accompanies PAM-mediated MBON21 inhibition (Fig. 6j), which likely puts the fly in a new spatial location that warrants reinvigorating foraging in response to the same sensory cue. Flexibly assigning values for the same action in accord with an animal’s own behavior may be advantageous in generating an efficacious foraging program that balances the potential gain and the associated energetic cost in accord with past efforts, current sensory contexts and current motivation. Finally, it is noteworthy that MBON21 receives KC inputs and these KC-MBON21 synapses might also be regulated by dopamine-induced plasticity just like KC-MBON09 synapses. Past experiences written by γ4 and γ5 DANs, like reward consumption, may play a role in regulating locomotor persistence through long-lasting alternation of KC-MBON21 synaptic weight. Future studies will reveal how this plasticity mechanism contributes to regulating locomotor parameters that may underlie optimal foraging strategies.

## ACKNOWLEDGMENTS

We thank Yoshinori Aso, Todd Roberts, Steven Shabel, Woj Wojtowicz, and members of the Hattori lab for their comments on the manuscript; Alexa De La Torre Schutz, Analia Marzoratti, Jack Mostyn and Julia Barasch for technical assistance and preliminary experiments; Peter Tsai, Wenhao Zhang for discussions; S. Lawrence Zipursky and Piero Sanfilippo for sharing flies and advice on Rdl visualization in MBON21; Gerald Rubin and Yoshinori Aso for sharing fly stocks before publication; and David Anderson, Vivek Jayaraman, Yulong Li, Bloomington Drosophila Stock Center (NIH P40OD018537) and Vienna Drosophila Resource Center for fly stocks. This work is supported by NIH R01DK132705.

## AUTHOR CONTRIBUTIONS

X.W., K.E.C. and C.C.S. performed experiments. X.W., K.E.C., C.C.S. and D.H. analyzed data.

K.E.C. designed 3D-printed parts for behavioral and imaging experiments. R.M. and A.M. setup hardware and software required for integration of fictrac. A.M. developed the MATLAB code for hemibrain connectome analysis and visualization. D.H. supervised the project. X.W. and D.H. wrote the manuscript with input from all authors.

## METHODS

### Fly Husbandry

For experiments that did not involve optogenetics, flies were reared in 25°C incubator with 12-hour/12-hour light/dark cycle. Within one day after eclosion, flies were genotyped under CO_2_ anesthesia and housed on regular food for 2-4 days. Approximately 24h before experiments, flies were transferred to new vials containing 1% agar for food deprivation, except for in Extended Data Fig. 2, where flies were transferred to regular food vials 24h before experiments (0h FD group) or agar vials 6h before experiments (6h FD group).

For experiments involving optogenetics, flies were reared in 25°C incubator in a dark box in vials with regular food until eclosion. Within one day after eclosion, flies were genotyped under CO_2_ anesthesia and housed in new vials containing regular food supplemented with all trans-Retinal (0.4mM) in 25°C incubator in a dark box for 2-4 days. Approximately 24h before experiments, flies were transferred to new vials containing 1% agar with 0.4mM all trans-Retinal for food deprivation.

For two-photon imaging experiments, tethered-fly behavioral experiments, and single-fly free-moving behavioral experiments, flies were housed as a pair of one male and one female. Female flies were used for two-photon imaging experiments. For tethered-fly behavioral experiments, a male or a female was used alternately for each experiment. For single-fly free-moving behavioral experiments, a male or a female was used alternately through separating the pair housed together into two vials without anesthesia. For multi-fly free-moving behavioral experiments, 2-3 males and 2-3 females were housed together and all flies were used for experiments (see below).

### Free-Moving Olfactory Behavior

The olfactory behavioral chamber (Extended Data Fig. 1a) comprises a 3D-printed resin frame with glass slides as the floor and ceiling that were sealed with UV glue (Bondic). Air flows were controlled by mass flow controllers (Aalborg, carrier stream 800mL/min, odor stream 200mL/min, vacuum 150mL/min) and odor was presented by switching the odor stream from a default port to an odor port connected to a custom mixing manifold through an 8-channel solenoid valve (Matrix). A custom MATLAB interface controlled the valve and mass flow controllers through a DAQ (National Instruments). Before each experiment, a flow meter was used to check whether the chamber was properly sealed when connecting the chamber to the vacuum line.

Video recording was performed using a camera above the chamber at 20 fps. Infrared lights were installed on both sides of the chamber for illumination and long-pass optical filters (Edmund Optics) were used to block LED light for optogenetic experiments. A custom MATLAB graphical user interface was used to control video recording and stimulus presentations, which issued timestamps for each trigger event. These timestamps were subsequently used to determine stimulus timings.

For optogenetic experiments, light stimulus was delivered from a strip of 655nm (for CsChrimson excitation) or 530nm (for GtACR1 excitation) high-power LEDs (Luxeon) positioned below the chamber that were controlled by an LED driver (BuckPuck). The light intensity measured at the chamber was 55.73μW/mm^2^ (655nm) and 49.38μW/mm^2^ (530nm).

For powdered-fly experiments, a single fly or 4–6 flies were introduced into a plastic vial with custom sift that contains a small amount of organic dye powder (reactive yellow 86, MP Biomedical). The vial was tapped to coat flies with the dye powder, and then flies were transferred into the arena using a foot pump. This procedure was performed with no anesthesia. For no-powder experiments, a pair of male and female were cold anesthetized and transferred into a clean arena. 10-15min of recovery time was given before experiments. Video recording and stimulus presentation were triggered manually when flies were grooming (for experiments with powdered or non-powdered flies) or standing still (for experiments with non-powdered flies) with minimum inter-trial intervals of >15s.

### Free-Moving Non-Olfactory Behavior

The non-olfactory behavioral chamber (Extended Data Fig. 1a) used for optogenetic stimulations is similar to the olfactory chamber except that it contains three arenas with no air inlet or outlet ports. Optogenetic light stimulus was delivered from a strip of 655nm LEDs positioned below the chamber. The light intensity measured at the chamber was 56.15μW/mm^2^ (high), 34.42μW/mm^2^ (medium), 10.47μW/mm^2^ (low). 4–6 flies were powdered and transferred into the arena using a foot pump. Video recording and stimulus presentation were triggered manually when most, if not all, flies were grooming.

### Two-Photon Functional Imaging

Two-photon functional imaging was performed using 2-photon microscope (Investigator, Bruker) equipped with ultra-fast laser (Chameleon Ultra2, Coherent). Images were obtained with 20x/1.0 NA water immersion objective (Olympus) typically using the following parameters: laser wavelength 925nm, laser power 9.6mW to 18mW after objective, pixel size 0.39 microns, pixel dwell time 4 microsec, and frame rate 18.2 fps. Details of imaging parameters different from above are listed in Extended Data Table 3.

Odors were delivered using a custom olfactometer with the same design described for free-moving olfactory behavior assays. Odor delivery line (carrier stream 800mL/min, odor stream 200mL/min) was split into two after mixing manifold with one directed to the fly and the other directed to the photoionization detector (mini-PID, Aurora Scientific) to monitor odor delivery for each trial. Odor was delivered through 1mm ID Teflon tubing, and the ends of the odor delivery line were connected to custom spouts with the end diameter of approximately 1/16 inch. The spout was placed in front of the fly head except for the experiments described in Extended Data Fig. 6f,g, in which the spout was placed on the left side of the fly head. PID signals were collected using the imaging software (PrairieView) at 100Hz.

Flies were prepared for functional imaging as described previously ^56^. Briefly, 3-to 5-day old females were cold anesthetized and head-fixed by the eyes using UV-cured glue (Bondic) to a metal shim (A-Laser) that was affixed to a custom 3D-printed saline reservoir. Proboscis was fixed in retracted position by UV glue. Head cuticle was removed in oxygenated saline (103 mM NaCl, 3 mM KCl, 5 mM TES, 26 mM NaHCO_3_, 1 mM NaH_2_PO_4_, 1.5 mM CaCl_2_, 4 mM MgCl_2_, 18 mM D(-) Ribose, adjusted to 270 - 275 mOsm bubbled with 95%O_2_/5%CO_2_, pH 7.2–7.3) using a 27-gauge × 1/2 hypodermic needle, and trachea and fat over the brain were removed using forceps. Oxygenated saline was perfused over the brain continuously during imaging at approximately 0.5 ml/min. Muscle 16 (frontal pulsatile organ) was pinched to minimize brain movement. After preparation, flies were placed on top of an air-supported ball in a dark chamber for recovery for at least 10 min before experiments.

Functional imaging experiments were performed with flies mounted on top of a resin (formlabs) ball (10 mm diameter, 0.13 g). The ball floats on 600mL/min air passing through 1mm diameter air canal of a 93 mm tall 3D-printed pedestal. Ball rotation was tracked using FicTrac software ^80^ using random patterns drawn on the ball surface with IR-absorbing ink (LDP).

Two orthogonally placed cameras (back view and side view of the fly; Flea3, FLIR) were used to aid fly mounting and to record fly behaviors. Pixel variance of fly leg movement was calculated from side-view video obtained at 20 fps, while ball rotation was tracked using FicTrac software from back-view video obtained at 60 fps. Optical filters (Edmund Optics) were installed before the cameras to pass IR light and block LED light used in optogenetic experiments. Cameras were controlled separately by custom MATLAB apps.

### Simultaneous Recording of Neural Responses and Alerting Behavior

To capture neural activity during alerting behavior in response to odors, ACV was delivered to the fly while she was standing still or grooming. Odor presentations were controlled manually when flies were grooming or standing still with a consistent inter-stimulus interval (ISI) that minimized sensory habituation. A custom MATLAB app was used to simultaneously trigger imaging and video recording of the fly. Video time was aligned to imaging time by identifying the onset and offset of two-photon laser captured in the videos.

Spontaneous locomotion of flies was captured under two conditions: “no air”, in which the air flow that delivers odor to the fly was turned off; “no odor” (PAM12 calcium recording), in which the air flow was on but no odor was presented.

### Functional Imaging with Optogenetic Stimuli

Flies were prepared for imaging as described above, but with visible light as dim as possible during the preparation. 655 nm (for CsChrimson excitation) or 530 nm (for GtACR1 excitation) ultra high power LED (Prizmatix) was used to deliver light. LED driver was controlled by the voltage-output function of the imaging software. Light intensity measured below the objective (power meter, Thorlabs PM16-130) was 409μW/mm^2^ for 655nm LED and 663μW/mm^2^ for 530nm LED. A custom MATLAB app was used to simultaneously trigger imaging, light protocol and odor protocol to allow the timing of light stimulation to be aligned with odor delivery.

### Tethered Fly Behavior

For tethered fly behavioral experiments, flies were prepared as described in two-photon imaging, except that head cuticle was intact and saline was not used. We noted an increased likelihood of locomotion during the baseline period before ACV. This is likely due to a general enhancement of activity level when flies were tethered without powder. For optogenetic experiments, ACV, ACV + light, or light trials were manually triggered while flies were standing still or grooming. A custom MATLAB app was used to simultaneously trigger PID recording, light protocol, and video recording of the fly.

### Immunostaining and Confocal Microscopy

Brain dissection, immunostaining, and confocal imaging were performed essentially as described in ^45^. Briefly, brains were dissected in 1xPBS, fixed for 1.5h at room temperature with 2% PFA in PBL, washed multiple times with 1x PBS containing 0.3% Triton X-100 (PBST), blocked with 10% normal goat serum diluted in PBST over 15 minutes at room temperature (RT), incubated in primary antibody mix at 4°C overnight, washed multiple times in PBST, and incubated in secondary antibody mix at 4°C overnight, or at RT for more than 3hr, before final washes with PBST. Primary antibodies used were rabbit anti-DsRed (1:1000, TaKaRa), mAb anti-bruchpilot (nc82, 1:10, DSHB), and chicken anti-GFP (1:1000, aves). Secondary antibodies were Alexa Fluor 568 goat anti-rabbit (1:200, Invitrogen), Alexa Fluor 633 goat anti-mouse (1:200, Invitrogen), and Alexa Fluor 488 goat anti-chicken (1:200, Invitrogen). Images were captured on a Zeiss LSM 880 using a 20X (0.8 NA) objective.

For Rdl visualization in MBON21, flies were heat-shocked in 37°C water bath for 3h at later third instar and early pupae stage and were dissected as 2-5d old adults. The same dissection and immunostaining protocols were employed as above except that dissection was performed in ice-cold Schneider’s insect medium (S0146, Sigma-Aldrich) and brains were incubated in primary and secondary antibodies for 2 overnights each. Primary antibodies were mouse IgG2a-anti-V5 (1:200 abcam cat. #ab27671) and rabbit anti-DsRed (1:500), and secondary antibodies were Alexa Fluor 488 goat anti-mouse (1:500) and Alexa Fluor 568 goat anti-rabbit (1:500).

### Quantification and statistical analysis

#### Two-Photon Imaging

Raw fluorescence signal was calculated as described in ^45^. Imaging frame were first registered across frames by a subpixel registration algorithm ^81^ using a visually verified baseline average image (typically 1-second before stimulus presentation) as a template, and the quality of registration was visually confirmed. Regions of interest (ROIs) were drawn manually based on the averaged image after registration, and mean pixel intensity within the ROI was extracted as raw fluorescence value representing signal in each frame. Raw fluorescence traces were converted into ΔF/F_0_ or Z-score traces using mean and standard deviation of a baseline period 1 seconds immediately before odor or light onset for odor/light presentation experiments, or 1 second immediately before movement onset/offset for experiments examining neural activity during spontaneous movement, except for Fig 6h,i, where 10th percentile was used as F_0_ to count for spontaneous firing of DANs. For imaging experiments without optogenetic stimulations, odor onset was determined by an inflection in PID signal, which was confirmed with manual inspection, or by using MATLAB-generated odor-trigger timestamps for those trials with no available PID data. For imaging experiments with optogenetic stimulations, odor onset was determined by MATLAB-generated timestamps and light onset was determined by voltage output protocol of the imaging software. If multiple trials of the same type for a fly, ΔF/F_0_ traces for these trials were averaged and generated one trace for the fly to use for statistical test, unless otherwise noted. For statistical analysis and comparisons between trial types and groups, ΔF/F_0_ traces were averaged or integrated over odor and/or light period. Flies were excluded from the analysis if the neuron recorded (KC calyx, DA sensor in MBON09) did not respond to any odor presentations during recording, with responding defined as Z-score reaching 2 within the first second of odor presentation and average Z-score during 2s odor above 2. Approximately 1-3 flies were eliminated from each group.

For KC somatic odor responses, three H2O trials (solvent) were included to distinguish ACV-responsive somas from solvent-responsive somas. Cells were selected if significantly responded to ACV at least in one trial but without significant responses in any H2O trials. Significant responses are defined as average Z-score during 2s odor period > 2.57 (equals to one tailed p < 0.005), using 2s immediately before odor as baseline. ΔF/F_0_ traces for cells from a fly were averaged to generate ΔF/F_0_ traces of the fly for each ACV trial. ΔF/F_0_ traces for each trial were averaged over ACV period, generating a neural response value to each ACV presentation for statistical tests.

To calculate the conditional probability of alerting given a particular MBON09 response amplitude, trials were separated into ten bins based on MBON09 response amplitudes with each bin containing 10% of the trials. This conditional probability of alerting given MBON09 response in each bin was then calculated as:

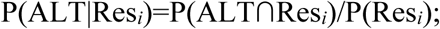

where P(Res*_i_*) is the probability of MBON09 response falling into bin *i* of 10 equally divided bins (i.e., 0.1) and P(ALT∩Res*_i_*) is the probability of co-occurrence of alerting and MBON09 response that falls into bin *i* (i.e., the number of alerting trials in bin *i* divided by the total number of trials). P(ALT|Res*_i_*) was plotted in Fig. 3 and Extended Data Fig. 5, with x axis values to be the center of each bin.

For ROC analysis, 2000 linearly increasing thresholds were generated between the minimum and maximum values of the predictor (MBON09 response or locomotion duration). For each threshold, trials were classified into two classes based on whether the predictor values were higher than the threshold. True positive rates and false positive rates were calculated comparing the classification with the ground truth (whether fly alerted in Fig. 3 and Extended Data Fig. 5, whether MBON21 activity was below 0 in Fig. 5, and whether CRE011 baseline activity was above 0 in Fig. 6) for each threshold. The observed area-under ROC curve values were compared with null distributions of area-under ROC curve values obtained from shuffled data (10000 iterations).

To compare PAM12 calcium activity or γ3 dopamine release across three behavioral contexts (alerting, non-alerting, spontaneous locomotion), the average Ca^2+^ activity or dopamine signal from 1s before starting locomotion till 5s after locomotion initiation was calculated for each fly and used to approximate locomotion-related PAM12 activity for the fly. The time of this ΔF/F_0_ trace was then aligned to alerting onsets (locomotion initiation after odor onsets) for each alerting trial, generating one ΔF/F_0_ trace of locomotion-related PAM12 activity corresponding with one alerting trial. To calculate the net response of odor and motion, fly’s average ΔF/F_0_ trace for non-alerting trials was added to those adjusted locomotion ΔF/F_0_ traces. Average ΔF/F_0_ over ACV period was calculated for each alerting trial and corresponding odor + motion trial for comparisons.

Spontaneous activity of PAM07/08 was determined by a sliding window method for inflection detection. The maximum raw fluorescence value in a 0.2s baseline window smaller than the minimum in the following 0.5s activity window was identified as an inflection (onset of Ca^2+^ increase). The inflection was counted as a real transient if average Z-score during the activity window is more than 2. Only one inflection is detected within 0.5s.

### Pixel Variance

Pixel variance of the selected ROI was calculated across a 9-(free-moving flies) or 7-(tethered flies) frame sliding window and used as locomotion proxy of the middle frame. This operation was done across the length of the movie to generate movement traces for the trial. For free-moving behavioral experiments, the ROI was the whole arena area, while for two-photon imaging and tethered fly behavioral experiments, the ROI was a rectangle area encompassing fly legs. The size of rectangle was consistent across experiments with the same field of view of the camera. Each experiment was performed with all relevant controls under the same illumination parameters.

For free-moving behavioral experiments, pixel variance was first normalized by dividing it with the number of flies in the arena. Then, Δ pixel variance was calculated by subtracting average pixel variance during baseline (0.5s before odor and/or light onset). If multiple trials of the same type existed for the same arena, Δ pixel variance traces for these trials were averaged and generated one trace for the arena to use for statistical test, unless otherwise noted. For free-moving multi-fly experiments without annotation (strawberry odor experiment), trials are omitted from analysis if average pixel variance during baseline (0.5s before odor onset) is above 2 to eliminate trials that flies were moving at odor onset. This pixel variance threshold is determined based on manual inspection of multiple videos. Less than 10% of trials are omitted. For two-photon imaging and tethered fly behavioral experiments, pixel variance is directly used to determine fly behavior. For statistical analysis and comparisons between trial types and groups, Δ pixel variance traces were averaged over 2.5s period since odor and/or light started.

### FicTrac

Frames that FicTrac failed to track and reset the position to 0 were teleported to the previous position, and missing frames were linearly interpolated. Fly x and y position and heading were gaussian smoothed with a window size of 60 (60 fps video), before calculating translational velocity (Euclidean norm of vector [x, y]), angular velocity (difference in heading), and net motion (Euclidean norm of vector [x, y, heading]). Data smoothing induces an artifactual inflection of motion preceding odor onset.

### Free-moving Fly Speed

For free-moving single fly experiment, the fly ROI was detected by subtracting the background image from each frame and the centroid of the ROI was used to calculate the speed of the fly. The x and y positions were first smoothed by a third-order median filter (20 fps video) before being used to calculate the translational velocity. Speed values above 139 mm/s were omitted and linearly interpreted using adjacent speed values.

### Annotation of Behavior

Frame-by-frame annotation of behaviors for free-moving flies were done as described previously^45^. A custom MATLAB GUI was made to allow annotation of multiple flies in the current video. In each frame, a fly was assigned with one of the three behavioral states, grooming with forelegs, grooming with hindlegs, or not grooming. Onset of grooming was assigned to the frame when the fly first exhibited movement immediately leading to grooming (e.g., when both forelegs were around fly head, or when both hindlegs were lifted from the ground). The offset of grooming was assigned to the frame that was one frame after the fly terminated grooming (e.g., when the fly separated its front legs from its head or from each other, or when both of the hindlegs touched the ground). In free-moving multi-fly behavioral experiments, flies were not annotated if they were blocked by other flies.

For behavior annotation of flies without powder, grooming onset and offset were defined as above. Locomotion was assigned to frames if the fly tapped the wall/top/bottom of the chamber with alternating forelegs. Other behaviors were defined as below:

Courtship: the male exhibiting behaviors within 1 fly length from the female, including orienting, chasing, foreleg tapping, licking, wing extension and attempted copulation. Proboscis extension: flies extending proboscis to touch floor without grooming or courtship.

Flailing: flies falling off the chamber wall onto their back and thrashing around in an attempt to right itself.

Jumping: flies translocating without alternating legs.

Standing-still was assigned to frames when the fly was not changed in location and not performing any behavior listed above. This included slight shuffling of feet or abdomen bending.

### Determination of Alerting, Locomotion Onset, and Locomotor Duration

For two-photon imaging and tethered fly behavioral experiments, fly movement states (i.e., moving vs non-moving) were classified by thresholding FicTrac parameters and/or pixel variance based on manual inspection of videos:

Net motion > 1mm/s & pixel variance > 10, or

Pixel variance > 15 (if no FicTrac; MBON09 recording with BZA).

Average FicTrac and/or pixel variance values were calculated during baseline (1s before odor onset) and odor period (2s) and compared with the thresholds to classify trials into alerting (non-moving during baseline and moving during odor) and non-alerting (non-moving during baseline and during odor). For tethered fly behavior, baseline period was 0.5s before odor onset.

Termination of locomotion was defined as the beginning of the first pause (1s below movement threshold) in each locomotor bout. The duration of locomotion was the time between the first moving frame during odor period to the termination of locomotion.

To determine the onset of spontaneous locomotion, pixel variance traces were linearly interpolated to align with net motion traces. Data were binarized using the threshold listed above and smoothed. The onset of sustained locomotion over 2s after a pause that lasted at least 1s was counted as initiation of spontaneous locomotion. The end of locomotion was defined as the beginning of the first 1s-pause after locomotion onset.

For free-moving behavioral experiments, moving vs non-moving were classified using a threshold of 1.5 mm/s. Trials were counted as alerting if average baseline motion (0.25s or 1s before odor onset) was lower than the threshold & flies moved for a total duration of 0.5s or more within 2.5s after odor onset. The offset of locomotion was defined as the beginning of the first pause (0.5s or 1s) after locomotion onset. Duration of locomotion for alerting trials was counted from the onset to the offset of locomotion, except for in Fig. 5j and Extended Data Fig. 9d, where the duration of locomotion was quantified as the total number of moving frames after odor onset divided by fps. For behavioral experiments with annotation, trials in which the fly was grooming at odor onset and transitioned to non-grooming for at least 1s within 2.5s after odor onset were classified as alerting trials.

### Determination of turning direction and upwind displacement

The angular velocity of the flies was used to determine turning direction in Extended Data Fig. 6e. Integral of angular velocity during 2s of ACV period was calculated and compared with a threshold range from-0.1 to 0.1 rad. A value within the threshold range was counted as non-moving, while a value beyond the range was counted as turning to the right (>0.1 rad) or turning to the left (<-0.1 rad). Upwind displacement during odor is defined as the distance along the airflow axis fly traveled during the odor period.

### Statistical Test

Nonparametric tests were used for statistic tests. For comparisons between two independent groups, Wilcoxon’s rank sum test was used. For paired data, Wilcoxon’s signed rank test was used. When comparing more than two groups, Kruskal-Wallis test was used, with post hoc bonferroni test for multiple comparisons. For categorical data, Fisher’s exact test was used. 95% bootstrap confidence interval in line plots was determined by computing mean or median of 10000 bootstrap samples.

---

**Extended Data Fig. 1:**
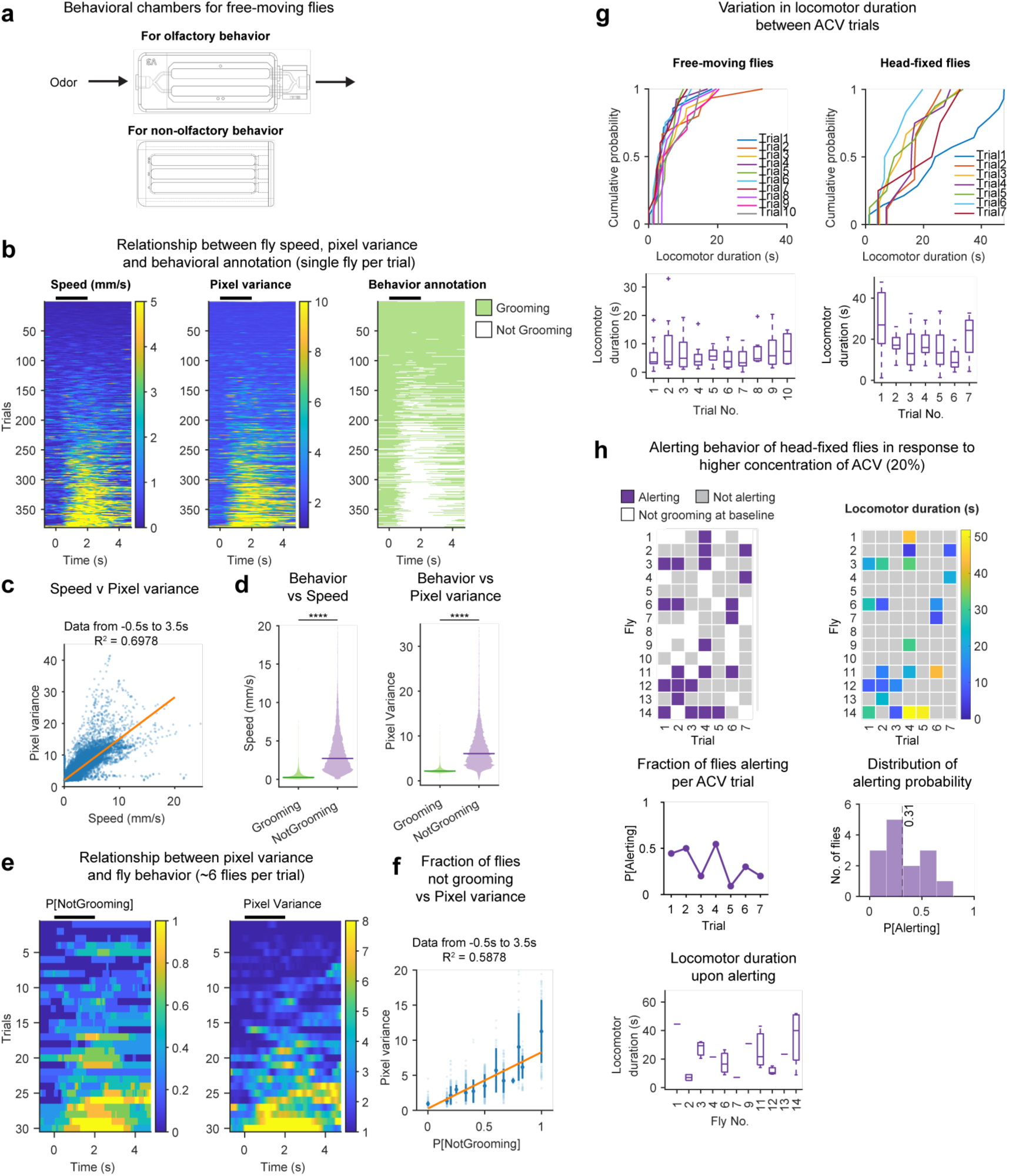
Analyses of alerting behavior. (a) Behavioral chambers for examining alerting behavior in free-moving flies. Top, 2-arena chamber for odor experiments; Bottom, 3-arena chamber for no-odor experiments. (b) Relationship between fly speed, pixel variance and behavioral annotation. Showing 380 trials with single fly in the chamber exposed to ACV. Black bars indicate odor period (same below). (c) Relationship between fly speed and pixel variance for each frame between-0.5s and 3.5s. N(frames)=30400; from 76 flies. (d) Relationship between behavioral annotation and fly speed/pixel variance. Left, comparison of fly speed in grooming vs non-grooming frames. Right, comparison of pixel variance in grooming vs non-grooming frames. Grooming frames, N=19382; non-grooming frames N=11018; from 76 flies. Solid lines indicate group median. Wilcoxon rank sum test. **** p<0.0001. (e) Relationship between fraction of flies not grooming and pixel variance. Showing 30 trials, each with 4–6 flies in the chamber exposed to ACV. (f) Relationship between fraction of flies not grooming (P[NotGrooming]) and pixel variance for each frame between-0.5s and 3.5s. Light blue dots indicate values for each frame. Dark blue dots indicate average pixel variance associated with given P[NotGrooming]; error bar, standard deviation. (g) Variation in locomotor duration across ACV trials. Top left, cumulative probability of moving duration for free-moving flies responding to 10 ACV trials. Bottom left, distribution of locomotion duration for free-moving flies responding to each ACV trial. Right, same as left for head-fixed flies. (h) Alerting behavior of food-deprived head-fixed flies in response to 20% ACV, as in Fig. 1.

**Extended Data Fig. 2:**
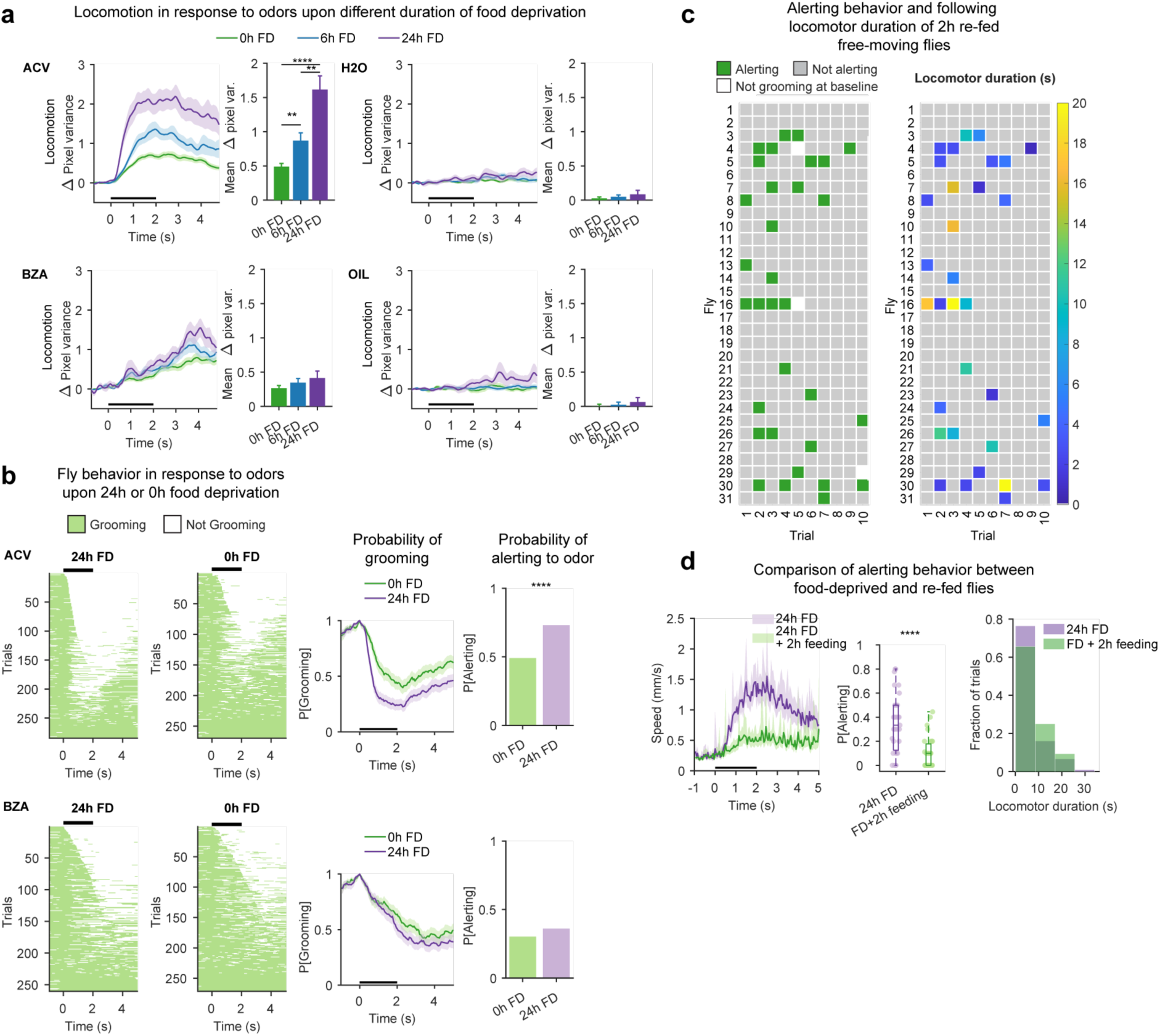
Starvation enhances alerting in response to ACV. (a) Alerting behavior of free-moving flies with 0h, 6h and 24h food deprivation (FD). BZA, Benzaldehyde. OIL, mineral oil (BZA solvent). Mean±SEM (same for all Δpixel variance line plots). N=66 (0h), 34 (6h), 32 (24h) experiments; 4–6 flies per experiment. Kruskal-Wallis test with post hoc Bonferroni test. **p<0.01; ****p<0.0001. (b) Fly behavior in response to ACV or BZA, with 0h or 24h food deprivation. Ethograms showing trials from 67 flies (0h) and 70 flies (24h). Line plots, mean±95% confidence interval (same for all P[Grooming] plots). Only trials in which the fly was grooming at odor onset are included. Fisher’s exact test. ****p=6.5E-9. (c) Alerting behavior of re-fed free-moving flies in response to ACV presentations. Left, trial-by-trial behavior classification; right, locomotor duration upon alerting. (d) Comparison of alerting behavior in response to ACV between food-deprived vs re-fed flies. Wilcoxon rank sum test. ****p=3.8E-5.

**Extended Data Fig. 3:**
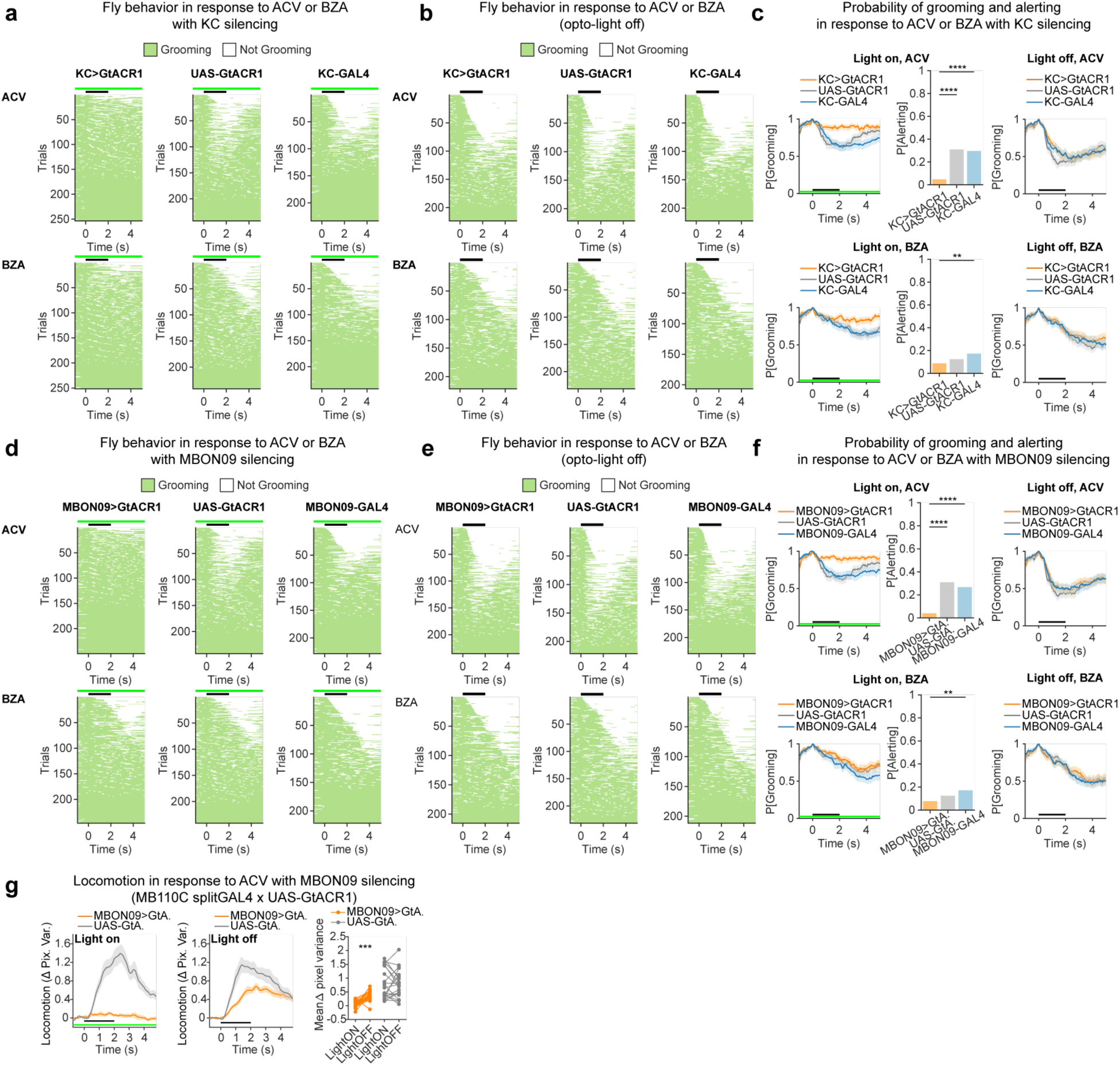
Effects of silencing KCs or MBON09 on alerting behavior. (a) Effect of KC silencing upon behavior in response to ACV or BZA (behavioral annotation for random subset of data in Fig. 2d). Only trials in which the fly was grooming at odor onset are included. Black bars indicate odor period and green bars indicate light period (same below). Trials from 59 (KC>GtA.), 55 (KC-GAL4), 54 (UAS-GtA.) flies. (b) Same as (a) for trials without light. (c) Probability of grooming over time and probability of alerting in response to ACV or BZA upon KC silencing. Fisher’s exact test. **p<0.01; ****p<0.0001. (d-f) Effect of MBON09 silencing upon behavior in response to ACV or BZA as in (a-c). Trials from 55 flies (MBON09>GtA.), 56 flies (MBON09-GAL4), 54 flies (UAS-GtA.). Note UAS-GtA flies are shared with data in (a). (g) Effect of MBON09 silencing upon behavior in response to ACV using a different driver (MB110C splitGAL4). Right panel, lines represent individual experiments with 4–6 flies per experiment. N= 20 (MBON09>GtA.) and 21 (UAS-GtA.) experiments. Wilcoxon signed-rank test. ***p=0.0009.

**Extended Data Fig. 4:**
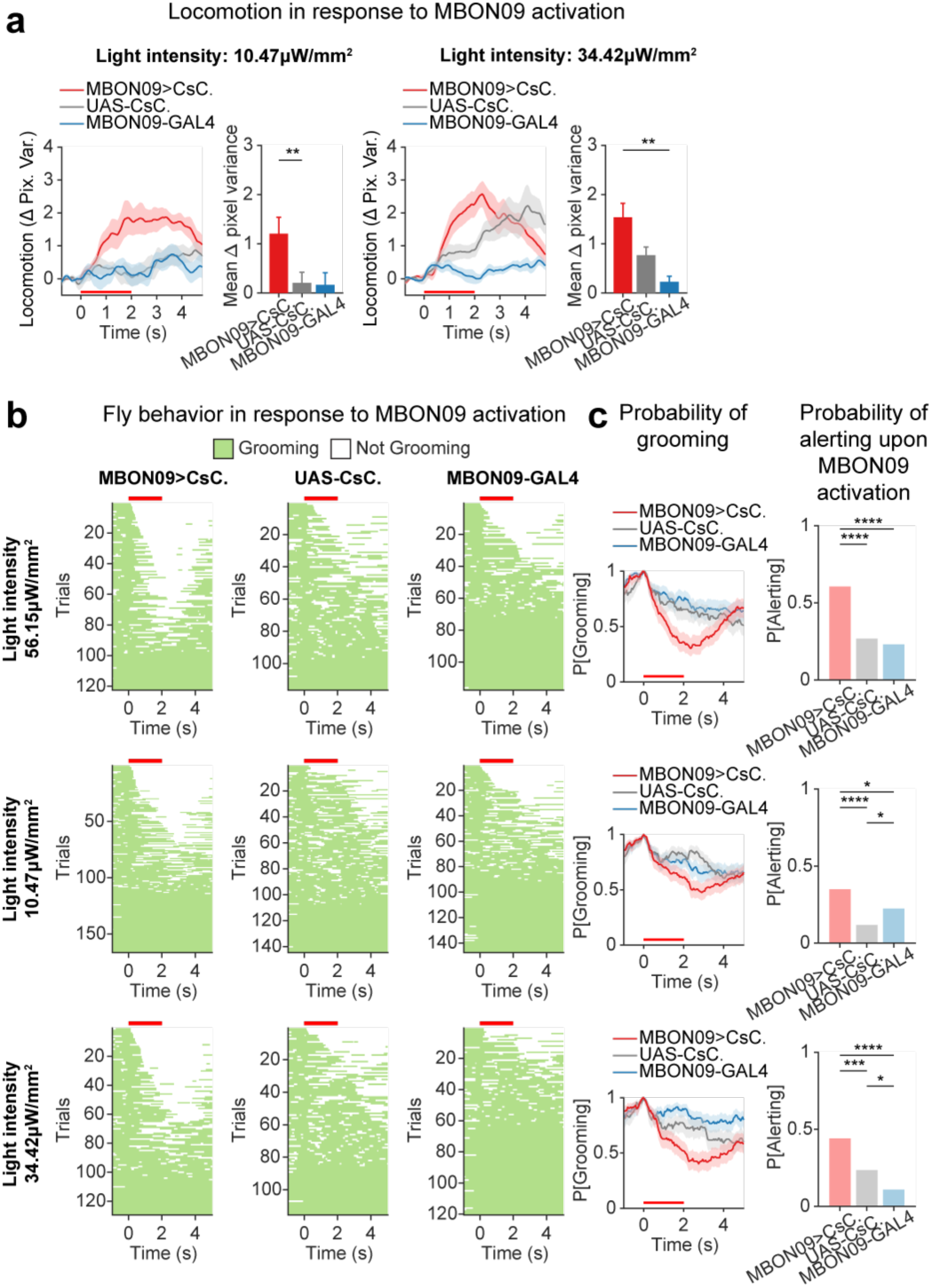
Effects of activation of MBON09 on alerting behavior. (a) Effect of optogenetic MBON09 activation upon locomotion. Lower light intensities than Fig. 2f. Red bars, light period. Data from 10 (MBON>CsC.), 10 (MBON-GAL4), 10 (UAS-CsC.) individual experiments with 4–6 flies per experiment. Kruskal-Wallis test with post hoc Bonferroni test. **p<0.01. (b) Effect of optogenetic MBON09 activation upon locomotion (behavioral annotation). Trials from 55 flies (MBON09>CsC.), 55 flies (MBON09-GAL4), 55 flies (UAS-CsC.). (c) Probability of grooming over time and probability of alerting in response to optogenetic MBON09 activation. Fisher’s exact test. *p<0.05; ***p<0.001; ****p<0.0001.

**Extended Data Fig. 5:**
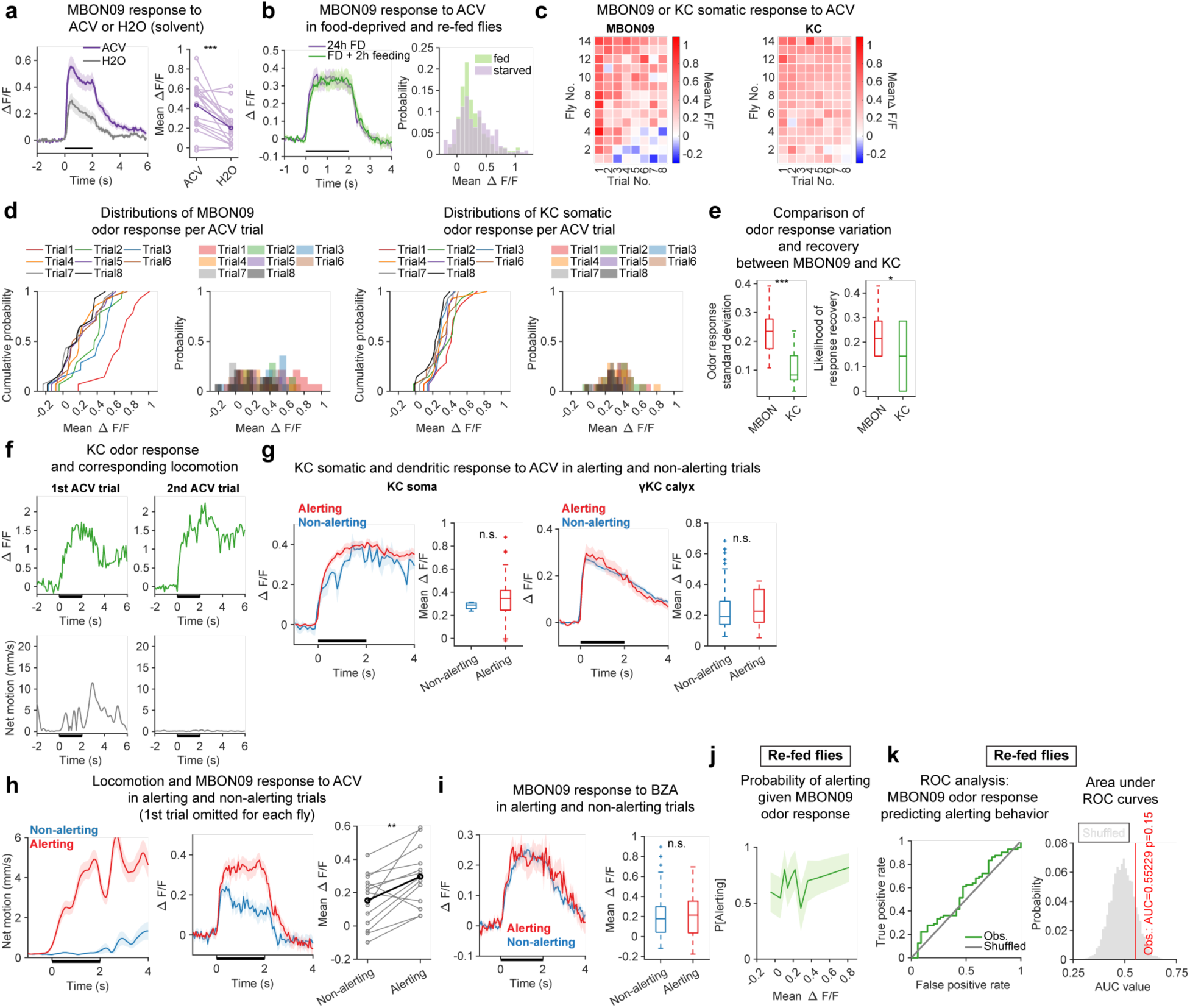
MBON09 and KC odor response and its association with alerting behavior in various contexts. (a) MBON09 calcium activity upon ACV or H2O (solvent) stimulations in food-deprived flies (measured at dendrites). Black bar indicates odor period (same below). Left, Mean±SEM (same for all imaging line plots). Right; light purple lines, flies; dark purple line, group average. N=18 flies. Wilcoxon signed-rank test. ***p=5.4E-4. (b) MBON09 ACV response (at soma) in food-deprived (FD) vs re-fed flies (FD then 2h feeding). Right, distributions of average MBON09 response during odor. N = 205 trials from 24 flies (food-deprived) and 125 trials from 15 flies (re-fed). (c) MBON09 and KC somatic response to ACV for eight trials in food-deprived flies. Mean ΔF/F during 2s odor period. For KCs, odor responses are averaged across multiple cells in the same fly (see Method). (d) Cumulative probability and distribution of odor response in eight ACV trials for MBON09 and KCs. (e) Comparison of odor response variation and recovery between MBON09 and KCs. Left, standard deviation of odor responses across trials per fly. Right, likelihood of odor response recovery. Odor response is considered recovered if ΔF/F observed in one trial is ≥0.1 higher than previous trial. Wilcoxon rank sum test. *p=0.027; ***p=0.00031. (f) Example KC calcium activity in response to two ACV presentations and corresponding fly locomotion in one KC soma. Trials are separated by ∼ 1 min. (g) KC somatic ACV response (left) and γKC calyx ACV response (right) in food-deprived flies in alerting vs non-alerting trials. N=14 (soma), 10 (calyx) flies. Wilcoxon rank sum test. p>0.05. (h) MBON09 ACV response in food-deprived flies in alerting vs non-alerting trials as in Fig. 3d, with first ACV trial for each fly omitted. Wilcoxon signed-rank test. **p=0.0081. (i) MBON09 BZA response in food-deprived flies in alerting vs non-alerting trials, as Fig. 3d,e. N=35 flies. Wilcoxon rank sum test. p>0.05. (j,k) Probability of alerting and ROC analysis for re-fed flies as in Fig. 3f,g.

**Extended Data Fig. 6:**
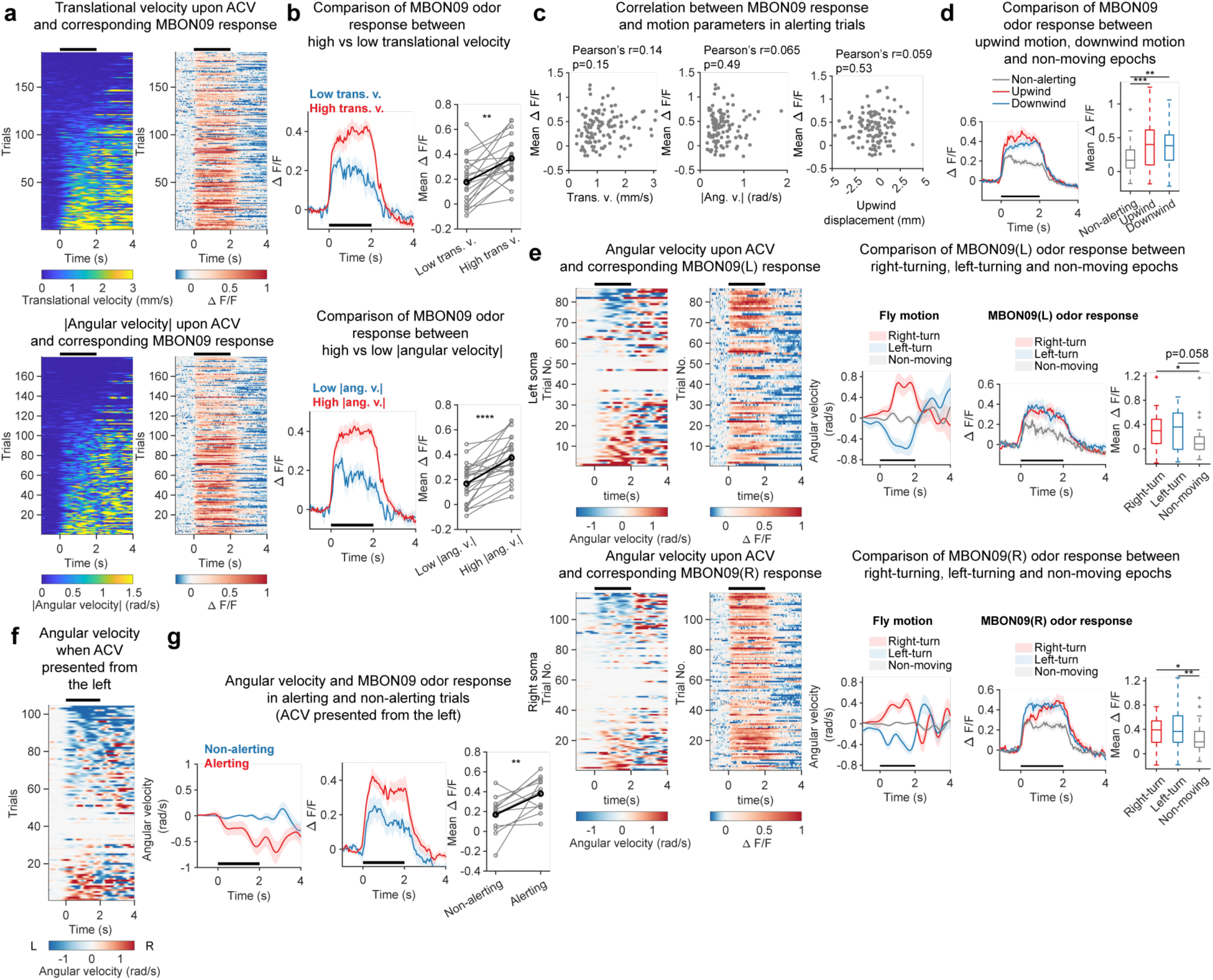
MBON09 odor response is associated with locomotor initiation regardless of moving direction. (a) Translational velocity (top) or magnitude of angular velocity (bottom) upon ACV presentation and corresponding MBON09 response in food-deprived flies. Sorted by behavior. Black bars indicate odor period (same below). N=186 trials (translational velocity) and 179 trials (|angular velocity|) from 24 flies. Trials in which fly was moving before odor onset are excluded. (b) MBON09 ACV response in food-deprived flies in high vs low translational velocity (trans. v.; top) or |angular velocity| (ang. v.; bottom) trials. Threshold for translational velocity, 0.5 mm/s; threshold for |angular velocity|, 0.1 rad/s. Flies that did not have both high and low trials are excluded. Wilcoxon signed-rank test. **p=0.003, ****p=7E-5. (c) Relationship between MBON09 ACV response and motion parameters in alerting trials. (d) MBON09 ACV response in upwind, downwind, and non-alerting trials. N=24 flies. Kruskal-Wallis test with post hoc Bonferroni test. **p=0.0013; ***p=7.02E-4. (e) Left (top) vs right (bottom) MBON09 ACV response in left-turn, right-turn, and non-alerting trials. Threshold for turning, >0.1 rad over 2s ACV period. Right turn, positive angular velocity. N=14 (right soma) and 10 (left soma) flies. Wilcoxon rank sum test. *p<0.05; **p<0.01. (f) Angular velocity in response to ACV that was presented from left side of fly. Right turn, positive angular velocity. Trials are excluded if flies move before odor presented. (g) Angular velocity and MBON09 ACV response of food-deprived flies in alerting vs non-alerting trials, when ACV was presented from left side. N=12 flies. Wilcoxon signed-rank test. **p=0.0093.

**Extended Data Fig. 7:**
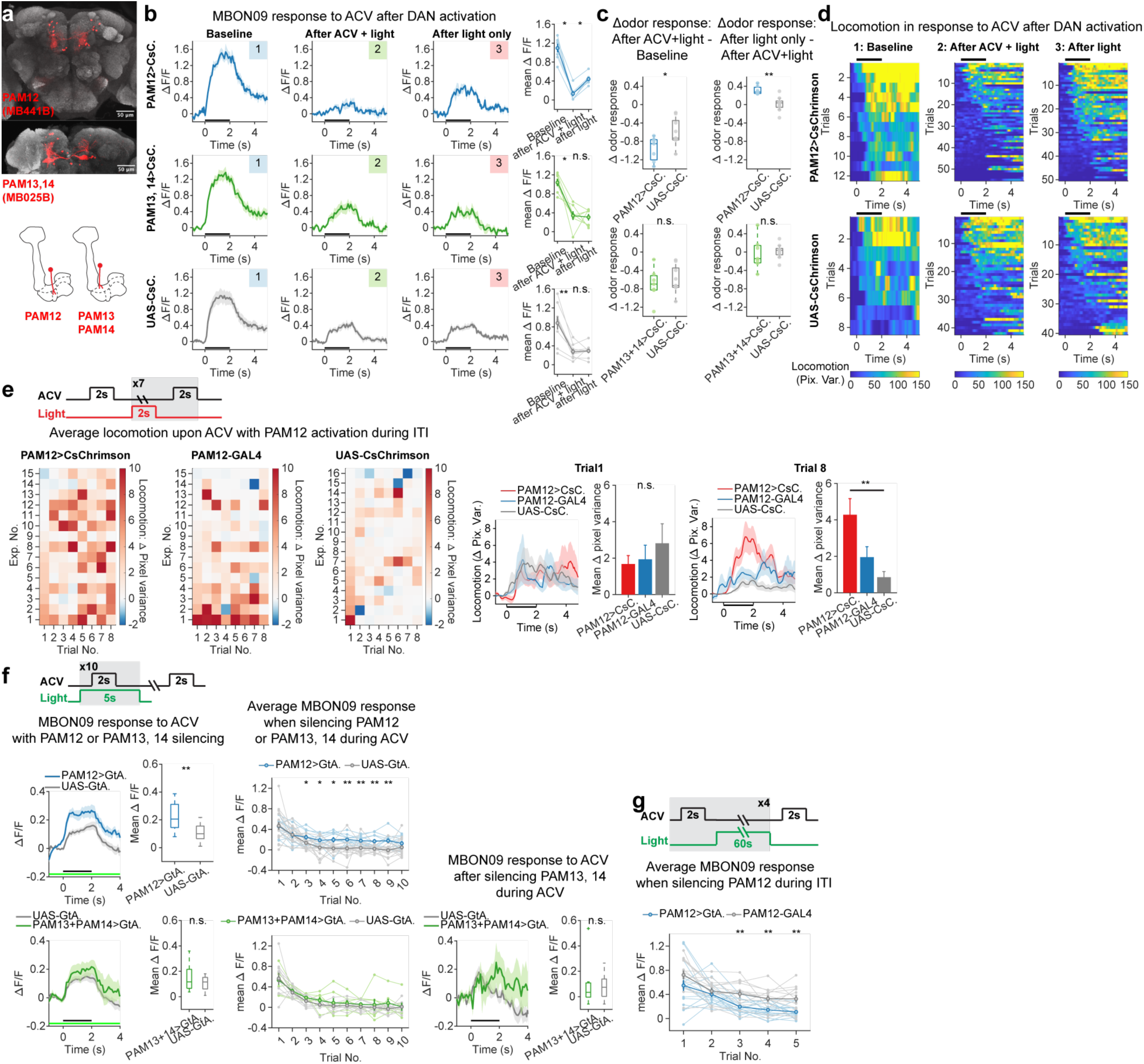
PAM12, but not PAM13/14, bidirectionally modulates MBON09 response and alerting behavior. (a) Confocal micrograph of brain showing expression pattern of MB441B split-GAL4 driver (PAM12, top) and MB025B split-GAL4 driver (PAM13 and PAM14, bottom). (b) MBON09 ACV response before and after optogenetic activation of PAM12 or PAM13/14 DANs with or without simultaneous presentation of ACV. Black bars, odor period (same below). Right, mean MBON09 ACV response before and after DAN activation. N=7 (PAM12>CsC.), 7 (PAM13-14>CsC.), 9 (UAS-CsC.) flies. Wilcoxon signed-rank test and rank sum test. *p<0.05; **p<0.01. (c) Changes in MBON09 ACV response upon ACV+DAN activation and upon DAN activation only for PAM12 (top) and PAM13/14 (bottom), as in Fig. 4d. Wilcoxon rank sum test. *p=0.031; **p=0.0021. (d) Head-fixed fly locomotion in response to ACV before and after optogenetic activation of PAM12 DANs. Trials are excluded if flies move before odor presentation. Note that baseline trials are the first ACV trial for each fly, while the other two types of trials were repeated three times for each fly. (e) ACV-induced locomotion of free-moving flies with 2s PAM12 activation during ITI. Up to 6 flies per experiment. Bottom, locomotion before activation of PAM12 (trial 1) and after activation of PAM12 during ITI for seven times (trial 8). Kruskal-Wallis test with post hoc Bonferroni; **p=0.0057. (f) MBON09 ACV response with PAM12 (top) or PAM13/14 (bottom) silencing during ACV. Light-shade lines represent individual flies whereas dark-shade lines represent group average. N=11 (PAM12>GtA.) and 17 (UAS-GtA.), and 7 (PAM13-14>GtA.) and 10 (UAS-GtA.) flies. Wilcoxon rank sum test. *p<0.05; **p<0.01. (g) Mean MBON09 ACV response with PAM12 silencing during ITI. Light-shade lines represent individual flies whereas dark-shade lines represent group average. N=17 (PAM12>GtA.), and 13 (PAM12-GAL4) flies. Wilcoxon rank sum test. **p<0.01.

**Extended Data Fig. 8:**
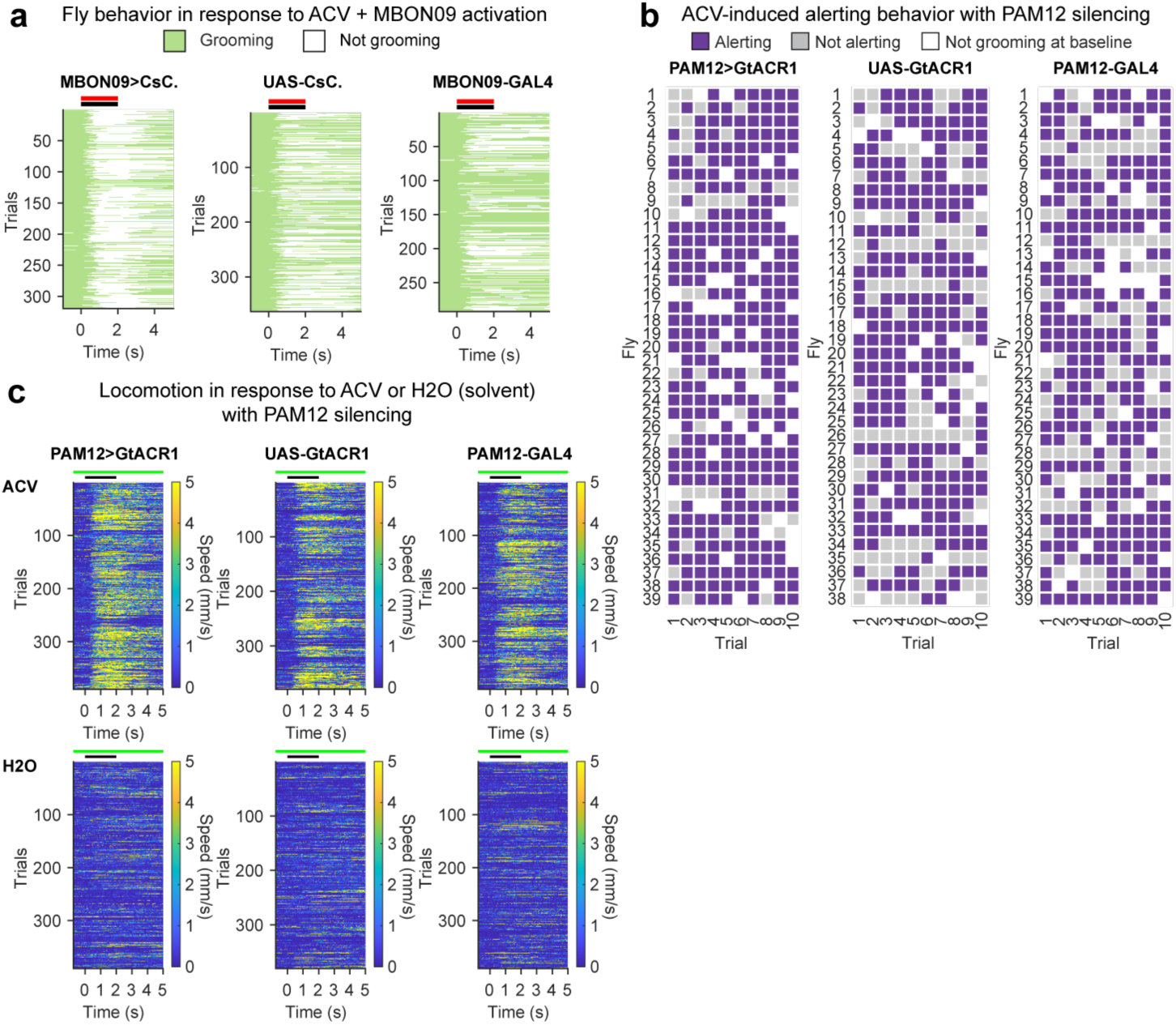
MBON09 activation and PAM12 silencing increase alerting consistence. (a) Effect of MBON09 optogenetic activation upon ACV-induced alerting (behavioral annotation). Red bar, light; black bar, ACV. Only trials in which fly was grooming at odor onset are included. 319 trials from 60 flies (MBON09>CsC.), 292 trials from 54 flies (MBON09-GAL4), 363 trials from 65 flies (UAS-CsC.). (b) Trial-by-trial classification of behavior in response to ACV in flies with PAM12 silencing. (c) Locomotion upon ACV and H2O (solvent) presentations in flies with PAM12 silencing. Green bar, light; black bar, ACV.

**Extended Data Fig. 9:**
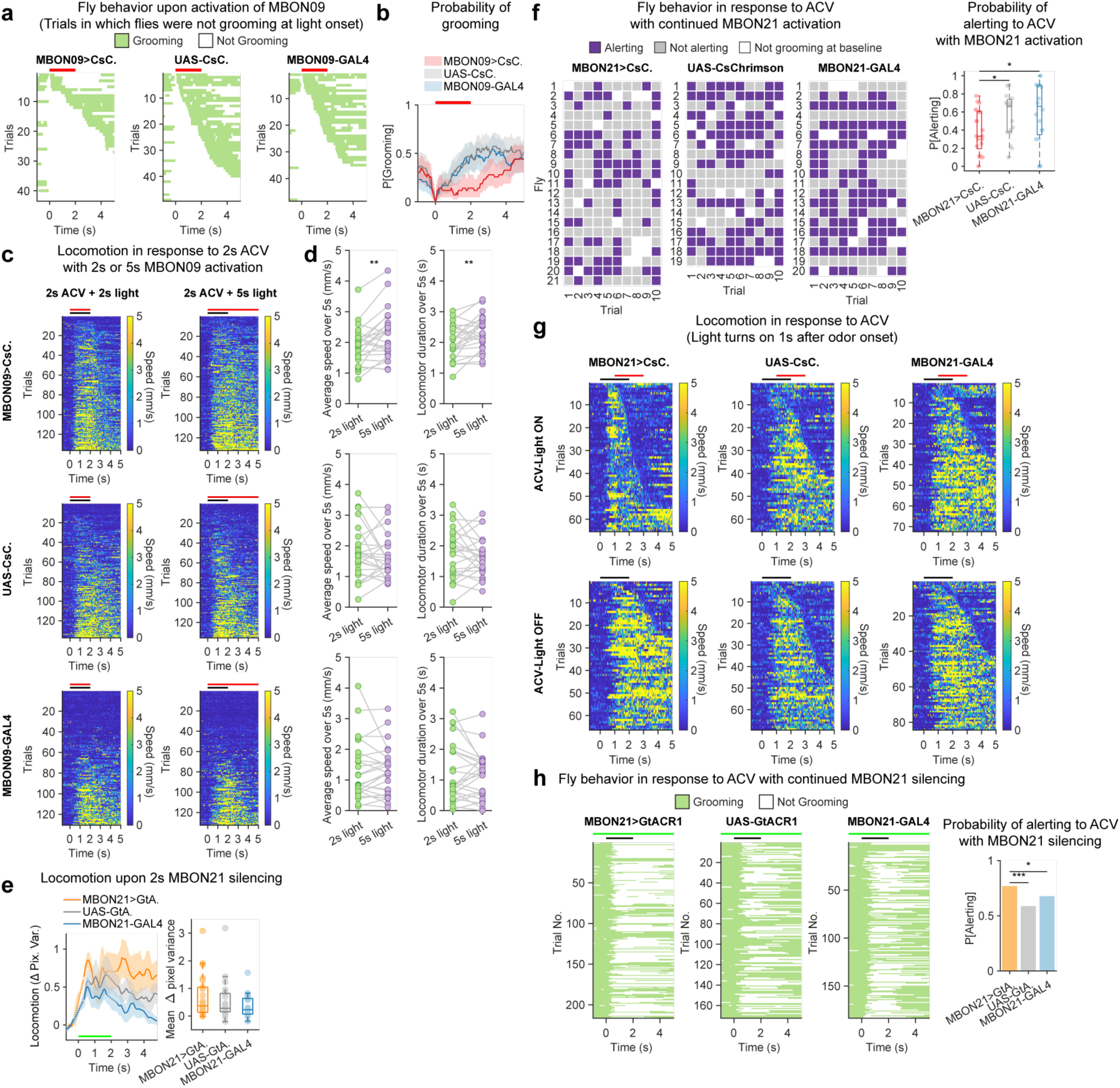
Effects of MBON09 activation and MBON21 manipulation on locomotor initiation and persistence. (a) Effect of optogenetic MBON09 activation on behavior when fly was not grooming at light onset. Red bars, light. Trials from 55 flies (MBON09>CsC.), 55 flies (MBON09-GAL4) and 55 flies (UAS-CsC.). (b) Effect of optogenetic MBON09 activation on probability of grooming when fly was not grooming at light onset. (c) Effect of 2s vs 5s optogenetic MBON09 activation on ACV-induced alerting. Black bars, ACV; red bars, light. Only trials in which fly was not moving before odor/light onset are shown. Trials from 24 (MBON09>CsC.), 24 (MBON21-GAL4) and 24 (UAS-CsC.) flies. (d) Effect of 2s vs 5s optogenetic MBON09 activation on mean speed or locomotor duration. Each line, individual fly. Wilcoxon signed rank test. **p<0.01. (e) Effect of acute optogenetic MBON21 silencing on locomotion. Green bar, light. Green-light-induced artifact on pixel variance was removed by linear interpolation and shown as dashed lines (left). N=22 (MBON21>GtA.), 22 (UAS-GtA.) and 20 (MBON21-GAL4) flies. Kruskal-Wallis test; p=0.45. (f) Effect of MBON21 activation upon ACV-induced alerting. Red light was continually on through 10 ACV trials. Left, trial-by-trial classification of alerting behavior. Right, probability of alerting per fly. Wilcoxon rank sum test; *p<0.05. (g) Effect of delayed MBON21 optogenetic activation upon locomotion in response to ACV. Only alerting trials are shown. N=22 (MBON21>CsC.), 17 (MBON21-GAL4) and 20 (UAS-CsC.) flies. (h) Effect of MBON21 silencing upon ACV-induced alerting. Green light was continually on through 10 ACV trials. N=26 (MBON21>GtA.), 19 (MBON21-GAL4), 20 (UAS-GtA.) flies. Fisher’s exact test. *p=0.0435; ***p=0.000167.

**Extended Data Fig. 10:**
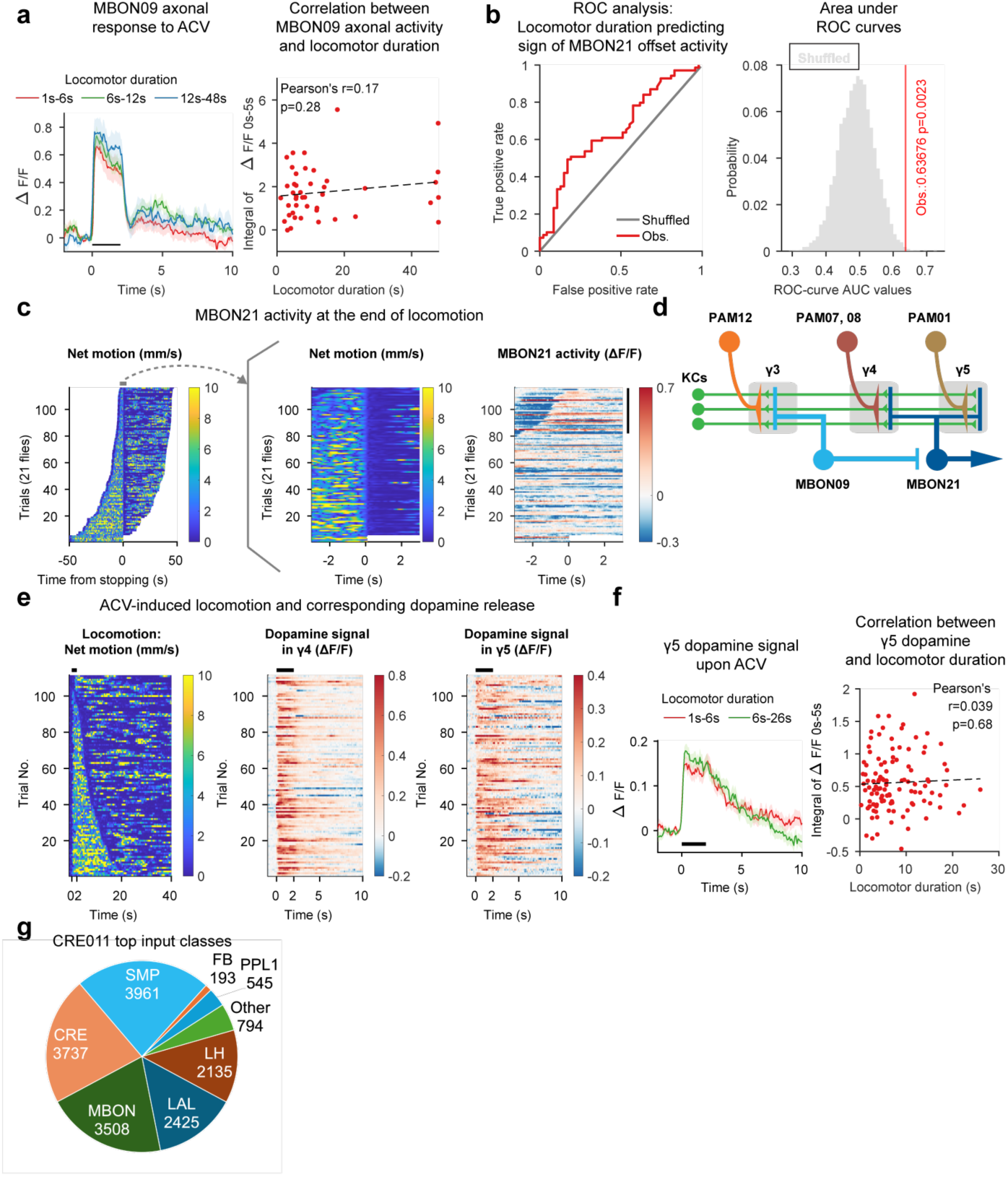
Relationship between locomotor duration/termination and MBON09/MBON21/γ4γ5 dopamine activity. (a) Relationship between MBON09 ACV response (axon) and resulting locomotor duration in alerting trials. N=14 (1-6 s), 15 (6-12 s) and 14 (12-48 s) trials from 11 flies. (b) Performance of locomotor duration as classifier of MBON21 offset (2.5-5s from ACV onset) activity sign as evaluated by ROC analysis. Left, ROC curves. Right, permutation test; red line indicates observed area under curve (AUC) value (obs.); gray histogram, distribution of AUC for shuffled data. One-tailed t-test. p=0.0023. (c) MBON21 activity at termination of ACV-induced locomotion. Left, locomotion in response to ACV in alerting trials, aligned to locomotor termination and sorted by locomotor duration. Center, zoomed-in locomotion. Right, corresponding MBON21 calcium activity. Vertical black bar indicates trials (trial 82-116) in which fly terminated locomotion within 3s after ACV offset, thus strong MBON21 inhibition in these trials represents ACV-induced inhibition. Trials from 21 flies. (d) Schematic of MBON09/MBON21 circuit. (e) Locomotion and corresponding dopamine release in γ4 and γ5 compartments in response to ACV presentations in alerting trials. Trials are sorted by locomotor duration upon alerting. Black bars, ACV (same below). Trials from 36 food-deprived flies. (f) Relationship between γ5 dopamine signal in response to ACV and resulting locomotor duration in alerting trials. N=56 (1-6 s) and 55 (6-26 s) trials from 36 flies. (g) CRE011 major (>100 connections) input neuron classes and numbers of connections. Other includes CL129, AVLP015, OA-VUMa7, PS166 and M_spPN4t9.

**Extended Data Table 1.**
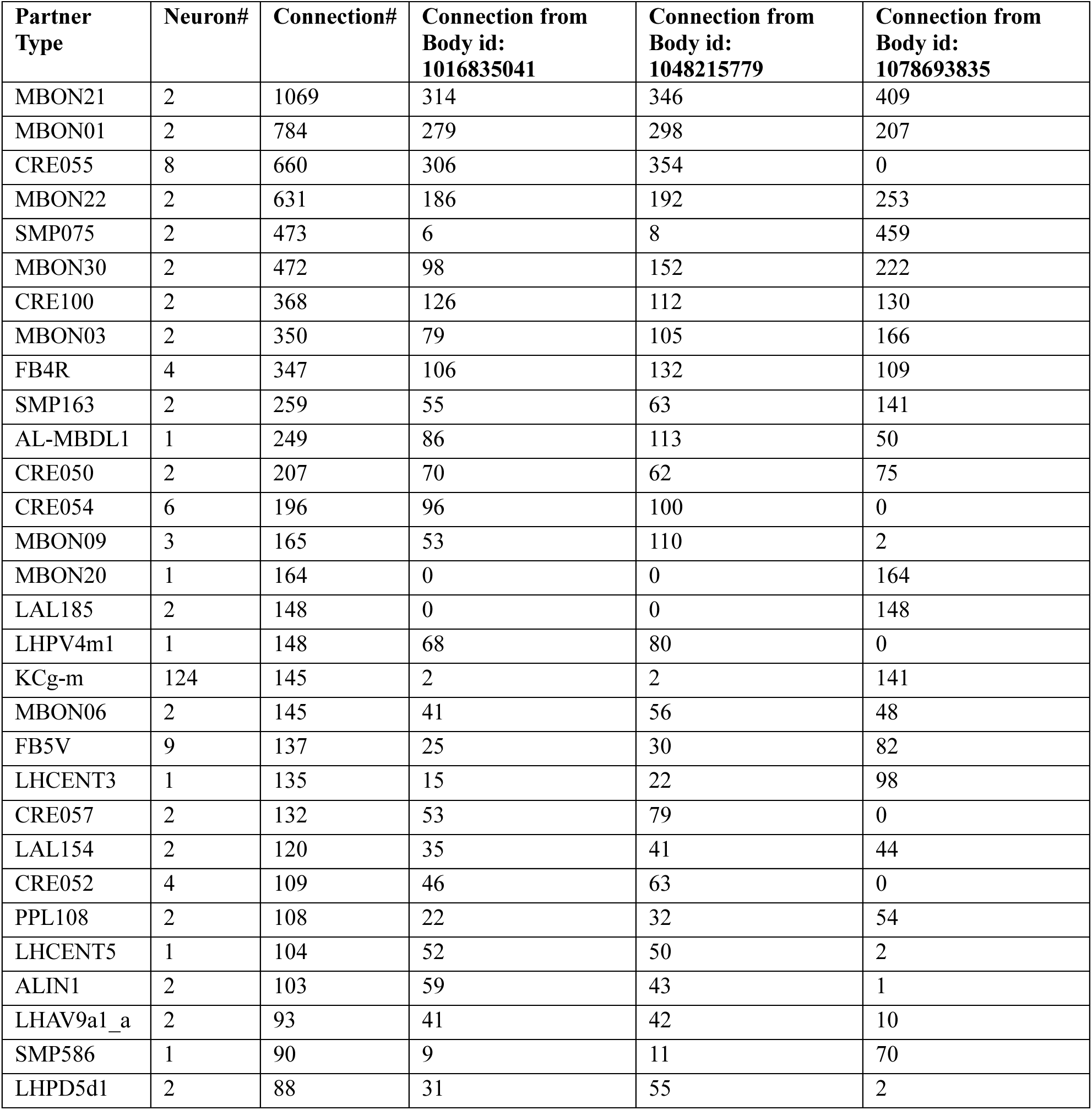
MBON09 top output connections.

| **Partner Type** | **Neuron#** | **Connection#** | **Connection from Body id: 1016835041** | **Connection from Body id: 1048215779** | **Connection from Body id: 1078693835** |
| --- | --- | --- | --- | --- | --- |
| MBON21 | 2 | 1069 | 314 | 346 | 409 |
| MBON01 | 2 | 784 | 279 | 298 | 207 |
| CRE055 | 8 | 660 | 306 | 354 | 0 |
| MBON22 | 2 | 631 | 186 | 192 | 253 |
| SMP075 | 2 | 473 | 6 | 8 | 459 |
| MBON30 | 2 | 472 | 98 | 152 | 222 |
| CRE100 | 2 | 368 | 126 | 112 | 130 |
| MBON03 | 2 | 350 | 79 | 105 | 166 |
| FB4R | 4 | 347 | 106 | 132 | 109 |
| SMP163 | 2 | 259 | 55 | 63 | 141 |
| AL-MBDL1 | 1 | 249 | 86 | 113 | 50 |
| CRE050 | 2 | 207 | 70 | 62 | 75 |
| CRE054 | 6 | 196 | 96 | 100 | 0 |
| MBON09 | 3 | 165 | 53 | 110 | 2 |
| MBON20 | 1 | 164 | 0 | 0 | 164 |
| LAL185 | 2 | 148 | 0 | 0 | 148 |
| LHPV4m1 | 1 | 148 | 68 | 80 | 0 |
| KCg-m | 124 | 145 | 2 | 2 | 141 |
| MBON06 | 2 | 145 | 41 | 56 | 48 |
| FB5V | 9 | 137 | 25 | 30 | 82 |
| LHCENT3 | 1 | 135 | 15 | 22 | 98 |
| CRE057 | 2 | 132 | 53 | 79 | 0 |
| LAL154 | 2 | 120 | 35 | 41 | 44 |
| CRE052 | 4 | 109 | 46 | 63 | 0 |
| PPL108 | 2 | 108 | 22 | 32 | 54 |
| LHCENT5 | 1 | 104 | 52 | 50 | 2 |
| ALIN1 | 2 | 103 | 59 | 43 | 1 |
| LHAV9a1_a | 2 | 93 | 41 | 42 | 10 |
| SMP586 | 1 | 90 | 9 | 11 | 70 |
| LHPD5d1 | 2 | 88 | 31 | 55 | 2 |

**Extended Data Table 2.** MBON21 top input connections.

| **Partner Type** | **Neuron#** | **Connection#** | **Connection to Body id: 5813022896** | **Connection to Body id: 985813153** | **Predicted neurotransmitter** |
| --- | --- | --- | --- | --- | --- |
| KCg-m | 589 | 7562 | 7551 | 11 | acetylcholine |
| MBON09 | 3 | 1069 | 503 | 566 | gaba |
| PAM07 | 12 | 584 | 307 | 277 | dopamine |
| KCg-d | 98 | 501 | 498 | 3 | dopamine |
| PAM08_b | 9 | 432 | 277 | 155 | dopamine |
| PAM08_c | 8 | 409 | 221 | 188 | dopamine |
| PAM08_a | 15 | 278 | 158 | 120 | dopamine |
| LAL185 | 2 | 275 | 271 | 4 | acetylcholine |
| DPM | 2 | 274 | 152 | 122 | dopamine |
| PAM01_a | 24 | 257 | 139 | 118 | dopamine |
| PAM01_b | 14 | 238 | 138 | 100 | dopamine |
| PAM08_e | 13 | 175 | 112 | 63 | dopamine |
| LHPV7c1 | 1 | 173 | 173 | 0 | acetylcholine |
| CRE030_b | 2 | 105 | 51 | 54 | glutamate |
| CRE024 | 2 | 72 | 52 | 20 | glutamate |
| KCg-t | 8 | 70 | 69 | 1 | dopamine |
| CRE031 | 4 | 63 | 44 | 19 | glutamate |
| PAM15_b | 3 | 60 | 39 | 21 | dopamine |
| SMP165 | 2 | 58 | 42 | 16 | glutamate |
| KCg-s2 | 1 | 52 | 52 | 0 | dopamine |
| PAM08_d | 2 | 50 | 30 | 20 | dopamine |
| CRE030_a | 2 | 48 | 46 | 2 | glutamate |
| MBON21 | 2 | 41 | 17 | 24 | acetylcholine |
| PAM12 | 19 | 39 | 24 | 15 | dopamine |
| MBON05 | 2 | 39 | 14 | 25 | glutamate |
| CRE029_a | 4 | 38 | 20 | 18 | glutamate |
| CRE059 | 4 | 36 | 31 | 5 | acetylcholine |
| KCg-s1 | 1 | 35 | 35 | 0 | dopamine |
| SMP079 | 2 | 34 | 25 | 9 | gaba |
| SMP127 | 2 | 32 | 15 | 17 | glutamate |

**Extended Data Table 3.** Additional 2-photon imaging settings.

| **Figure** | **Experiment description** | **Objective** | **Pixel size (microns)** | **Frame rate (fps)** |
| --- | --- | --- | --- | --- |
| **Fig. 5l; Extended Data Fig. 10e,f** | $\gamma$ lobe dopamine recording | 20x/1.0 NA water immersion objective (Olympus) | 0.47 | 10.9 |
| **Extended Data Fig. 5a** | MBON09 dendritic recording | 60x/1.0 NA water immersion objective (Olympus) | 0.13 | 18 |
| **Extended Data Fig. 5c- g (left)** | KC somatic recording | 20x/1.0 NA water immersion objective (Olympus) | 0.58 | 9.7 |
| **Extended Data Fig. 5g (right)** | KC calyx Ca<sup>2+</sup> recording | 20x/1.0 NA water immersion objective (Olympus) | 0.47 | 10.9 |

**Extended Data Table 4.** Sample size and statistics.

| Figure | Sample size | Data excluded | Statistical test | p value |
| --- | --- | --- | --- | --- |
| **Extended Data Fig. 1d** | Frame#<br>Grooming: 19382<br>Non-grooming: 11018<br>from 76 flies | None | Wilcoxon rank sum | Speed: 0<br>Pixel variance: 0 |
| **Extended Data Fig. 2a** | ROI# (4–6 flies/ROI)<br>0h FD: 66<br>6h FD: 34<br>24h FD: 32 | None | kruskal wallis | 9.97E-11 (ACV)<br>0.151(H2O)<br>0.206 (BZA)<br>0.408 (Oil) |
|  |  |  | Post hoc bonferroni | ACV: 0.0049 (0h vs 6h)<br>5.56E-11 (0h vs 24h)<br>0.0045 (6h vs 24h)<br>H2O: 1 (0h vs 6h)<br>0.2 (0h vs 24h)<br>-0.33 (6h vs 24h)<br>BZA: 0.95 (0h vs 6h)<br>0.25 (0h vs 24h)<br>1 (6h vs 24h)<br>Oil: 1 (0h vs 6h)<br>0.56 (0h vs 24h)<br>1 (6h vs 24h) |
| **Extended Data Fig. 2b** | Trial#<br>ACV:<br>0h FD: 269<br>24h FD: 284<br>BZA:<br>0h FD: 271<br>24h FD: 260<br>Fly #<br>0h FD: 67<br>24h FD: 70 | Baseline non-grooming trials (see Methods) | Fisher's exact test | 6.51E-9 (ACV)<br>0.167 (BZA) |
| **Extended Data Fig. 2d** | Fly#<br>24h FD: 31<br>24h FD +2h feeding: 31 | Baseline moving trials (see Methods) | Wilcoxon rank sum | 3.76E-5 |
| **Fig. 2d** | ROI# (4–6 flies/ROI)<br>KC>GtA: 35<br>KC-GAL4: 16<br>UAS-GtA: 56 | None | kruskal wallis | 1.46E-12 (ACV-Light ON)<br>0.00848 (BZA-Light ON)<br>0.714 (ACV-Light OFF)<br>0.932 (BZA-Light OFF) |
|  |  |  | Post hoc bonferroni | ACV-Light ON: 9.03E-12 (KC>GtA vs UAS-GtA)<br>4.51E-7 (KC>GtA vs KC-GAL4)<br>0.954 (UAS-GtA vs KC-GAL4)<br>BZA-Light ON: 0.018 (KC>GtA vs UAS-GtA)<br>0.032 (KC>GtA vs KC-GAL4)<br>0.815 (UAS-GtA vs KC-GAL4)<br>ACV-Light OFF: 0.993 (KC>GtA vs UAS-GtA)<br>0.776 (KC>GtA vs KC-GAL4)<br>0.696 (UAS-GtA vs KC-GAL4) |
|  |  |  |  | BZA-Light OFF: 0.961 (KC>GtA vs UAS-GtA)<br>0.992 (KC>GtA vs KC-GAL4)<br>0.942 (UAS-GtA vs KC-GAL4) |
| **Fig. 2f;**<br>**Extended**<br>**Data Fig.**<br>**4a** | ROI# (4–6<br>flies/ROI)<br>MBON09>CsC: 10<br>UAS-CsC: 10<br>MBON09-GAL4:<br>10 | None | kruskal<br>wallis | 0.0215 (Low intensity)<br>0.00156 (Medium intensity)<br>0.000572 (High intensity) |
|  |  |  | Post hoc<br>bonferroni | Low:<br>0.0509 (MBON09>CsC vs MBON09-GAL4)<br>0.0475 (MBON09>CsC vs UAS-CsC)<br>1 (MBON09-GAL4 vs UAS-CsC)<br>Medium:<br>0.00103 (MBON09>CsC vs MBON09-GAL4)<br>0.383 (MBON09>CsC vs UAS-CsC)<br>0.119 (MBON09-GAL4 vs UAS-CsC)<br>High:<br>0.00103 (MBON09>CsC vs MBON09-GAL4)<br>0.00691 (MBON09>CsC vs UAS-CsC)<br>1 (MBON09-GAL4 vs UAS-CsC) |
| **Fig. 2g** | ROI# (4–6<br>flies/ROI)<br>MBON09>GtA: 27<br>UAS-GtA: 56<br>MBON09-GAL4:<br>15 | None | kruskal<br>wallis | 1.26E-11 (ACV-Light ON)<br>0.311 (BZA-Light ON)<br>0.336 (ACV-Light OFF)<br>0.078 (BZA-Light OFF) |
|  |  |  | Post hoc<br>bonferroni | ACV-Light ON: 9.98E-12 (MBON09>GtA vs<br>UAS-GtA)<br>1.34E-5 (MBON09>GtA vs MBON09-GAL4)<br>0.86 (UAS-GtA vs MBON09-GAL4)<br>BZA-Light ON: 0.337 (MBON09>GtA vs<br>UAS-GtA)<br>0.447 (MBON09>GtA vs MBON09-GAL4)<br>0.977 (UAS-GtA vs MBON09-GAL4)<br>ACV-Light OFF: 0.321 (MBON09>GtA vs<br>UAS-GtA)<br>0.581 (MBON09>GtA vs MBON09-GAL4)<br>0.998 (UAS-GtA vs MBON09-GAL4)<br>BZA-Light OFF: 0.186 (MBON09>GtA vs<br>UAS-GtA)<br>0.0895 (MBON09>GtA vs MBON09-GAL4)<br>0.63 (UAS-GtA vs MBON09-GAL4) |
| **Fig. 2i** | ROI# (4–6<br>flies/ROI)<br>MBON09>GtA: 19<br>UAS-GtA: 18<br>MBON09-GAL4:<br>19 | Baseline<br>moving trials<br>(see<br>Methods) | kruskal<br>wallis | 3.11E-5 (Strawberry – Light ON)<br>0.158 (Strawberry – Light OFF) |
|  |  |  | Post hoc<br>bonferroni | Strawberry – Light ON: 1.72E-5<br>(MBON09>GtA vs UAS-GtA)<br>0.0275 (MBON09>GtA vs MBON09-GAL4)<br>0.148 (UAS-GtA vs MBON09-GAL4)<br>Strawberry – Light OFF: 0.985 (MBON09>GtA<br>vs UAS-GtA)<br>0.165 (MBON09>GtA vs MBON09-GAL4)<br>1 (UAS-GtA vs MBON09-GAL4) |
| **Fig. 2h** | Trial#<br>“Still” baseline:<br>MBON09>GtA:105<br>MBON09-GAL4:76<br>UAS-GtA:139<br>“Grooming” baseline:<br>MBON09>GtA:100<br>MBON09-GAL4:59<br>UAS-GtA:83<br>Fly#<br>MBON09>GtA:50<br>MBON09-GAL4:30<br>UAS-GtA:50 | Baseline non-grooming or non-still trials | Fisher’s exact test | “Still” baseline:<br>4.18E-15 (MBON09>GtA vs MBON09-GAL4)<br>2.4E-16(MBON09>GtA vs UAS-GtA)<br>0.465 (MBON09-GAL4 vs UAS-GtA)<br>“Grooming” baseline:<br>1.65E-21 (MBON09>GtA vs MBON09-GAL4)<br>3.6E-24(MBON09>GtA vs UAS-GtA)<br>0.842 (MBON09-GAL4 vs UAS-GtA) |
| **Extended Data Fig. 3c** | Trial#<br>ACV – Light on:<br>KC>GtA: 253<br>KC-GAL4: 233<br>UAS-GtA: 242<br>BZA – Light on:<br>KC>GtA: 251<br>KC-GAL4: 242<br>UAS-GtA: 241<br>Fly #<br>KC>GtA: 59<br>KC-GAL4: 55<br>UAS-GtA: 54 | Baseline non-grooming trials | Fisher’s exact test | ACV – Light on: 2.67E-15 (KC>GtA vs UAS-GtA)<br>5.54E-14 (KC>GtA vs KC-GAL4)<br>0.765 (UAS-GtA vs KC-GAL4)<br>BZA – Light on: 0.191 (KC>GtA vs UAS-GtA)<br>0.00487 (KC>GtA vs KC-GAL4)<br>0.16 (UAS-GtA vs KC-GAL4) |
| **Extended Data Fig. 3f** | Trial#<br>ACV – Light on:<br>MBON09>GtA:249<br>MBON09-GAL4:228<br>UAS-GtA:242<br>BZA – Light on:<br>MBON09>GtA:246<br>MBON09-GAL4:221<br>UAS-GtA:241<br>Fly#<br>MBON09>GtA:55<br>MBON09-GAL4:56<br>UAS-GtA:54 | Baseline non-grooming trials | Fisher’s exact test | ACV – Light on: 8.424E-13(MBON09>GtA vs MBON09-GAL4)<br>3.292E-16 (MBON09>GtA vs UAS-GtA)<br>0.36 (MBON09-GAL4 vs UAS-GtA)<br>BZA – Light on: 0.00269 (MBON09>GtA vs MBON09-GAL4)<br>0.0976(MBON09>GtA vs UAS-GtA)<br>0.188 (MBON09-GAL4 vs UAS-GtA) |
| **Extended Data Fig. 4c** | Trial#<br>Low intensity:<br>MBON09>CsC: 166<br>MBON09-GAL4: 147<br>UAS-CsC: 144<br>Medium intensity:<br>MBON09>CsC: 129 | Baseline non-grooming trials | Fisher’s exact test | Low:<br>0.0178 (MBON09>CsC vs MBON09-GAL4)<br>2.32E-6 (MBON09>CsC vs UAS-CsC)<br>0.0195 (MBON09-GAL4 vs UAS-CsC)<br>Medium:<br>3.29E-9 (MBON09>CsC vs MBON09-GAL4)<br>0.000742 (MBON09>CsC vs UAS-CsC)<br>0.0144 (MBON09-GAL4 vs UAS-CsC)<br>High:<br>5.27E-9 (MBON09>CsC vs MBON09-GAL4)<br>1.45E-7 (MBON09>CsC vs UAS-CsC) |
|  | MBON09-GAL4: 120<br>UAS-CsC: 115<br>High intensity: MBON09>CsC: 122<br>MBON09-GAL4: 117<br>UAS-CsC: 116<br>Fly#<br>MBON09>CsC: 55<br>MBON09-GAL4: 55<br>UAS-CsC: 55 |  |  | 0.547 (MBON09-GAL4 vs UAS-CsC) |
| **Extended Data Fig. 3g** | ROI# (4–6 flies/ROI)<br>MBON09>GtA: 20<br>UAS-GtA: 21 | None | Wilcoxon signed rank | ACV light on vs off:<br>0.00089 (MBON09>GtA)<br>0.689 (UAS-GtA) |
| **Fig. 3e** | Fly# 13<br>63 alerting trials<br>52 non-alerting trials | Flies without both alerting and non-alerting trials; baseline moving trials | Wilcoxon signed rank | 0.0081 |
| **Fig. 3h** | Trial#<br>Locomotion initiation: 43<br>Non-locomotion: 43<br>from 13 flies | None |  |  |
| **Fig. 3i** | Fly# 10<br>72 alerting trials<br>34 non-alerting trials | Flies without both alerting and non-alerting trials; baseline moving trials | Wilcoxon signed rank | 0.62 |
| **Extended Data Fig. 5a** | Fly# 18 | None | Wilcoxon signed rank | 5.36E-4 |
| **Extended Data Fig. 5e** | Fly #<br>MBON09: 14<br>KCs: 14 | None | Wilcoxon rank sum | 0.00031 (standard deviation)<br>0.027 (recovery rate) |
| **Extended Data Fig. 5g (left)** | Trial#<br>Alerting: 89<br>Non-alerting: 4<br>from 14 flies | Baseline moving trials | Wilcoxon rank sum | 0.36 (soma) |
| **Extended Data Fig. 5g (right)** | Trial#<br>Calyx:<br>Alerting: 15<br>Non-alerting: 89<br>from 10 flies | Baseline moving trials | Wilcoxon rank sum | 0.45 (Calyx) |
| **Extended Data Fig. 5h** | Fly# 13<br>52 alerting trials | Flies without both alerting and non- | Wilcoxon signed rank | 0.0081 |
|  | 49 non-alerting trials | alerting trials; baseline moving trials; 1<sup>st</sup> ACV trial |  |  |
| **Extended Data Fig. 5i** | Trial#<br>Alerting: 51<br>Non-alerting: 145<br>from 35 flies | Baseline moving trials | Wilcoxon rank sum | 0.73 |
| **Extended Data Fig. 6b** | Fly#<br>Translational velocity: 21<br>Angular velocity: 22 | Flies without both alerting and non-alerting trials; baseline moving trials | Wilcoxon signed rank | 0.003 (Translational velocity)<br>7E-5 (Angular velocity) |
| **Extended Data Fig. 6d** | Trial#<br>Non-alerting: 77<br>Upwind: 45<br>Downwind: 68<br>from 24 flies | Baseline moving trials | kruskal wallis | 1.08E-4 |
|  |  |  | Post hoc bonferroni | 0.0013 (Non-alerting vs Downwind)<br>7.02E-4 (Non-alerting vs Upwind)<br>1 (Upwind vs Downwind) |
| **Extended Data Fig. 6e** | Trials#<br>Right soma:<br>Right-turn: 32<br>Left-turn: 39<br>Non-moving: 47<br>from 14 flies<br>Left soma:<br>Right-turn: 31<br>Left-turn: 28<br>Non-moving: 28<br>from 10 flies | None | Wilcoxon rank sum | Right soma:<br>0.84 (Right-turn vs Left-turn)<br>0.0177 (Right-turn vs non-moving)<br>0.0025 (Left-turn vs non-moving)<br>Left soma:<br>0.779 (Right-turn vs Left-turn)<br>0.0131 (Right-turn vs non-moving)<br>0.0584 (Left-turn vs non-moving) |
| **Extended Data Fig. 6g** | Fly# 12 | Flies without both alerting and non-alerting trials; baseline moving trials | Wilcoxon signed rank | 0.0093 |
| **Fig. 4b** | Fly#<br>DANs>CsC: 8<br>DANs-GAL4: 9 | None | Wilcoxon rank sum | 0.54 |
| **Fig. 4c** | Fly#<br>DANs>CsC: 8<br>DANs-GAL4: 9 | None | Wilcoxon signed rank | DANs>CsC:<br>0.0078 (Baseline vs After ACV+light)<br>0.0078 (After ACV+light vs After light)<br>DANs no CsC:<br>0.426 (Baseline vs after ACV+light)<br>0.82 (After ACV+light vs After light) |
| **Fig. 4d** | Fly#<br>DANs>CsC: 8<br>DANs-GAL4: 9 | None | Wilcoxon rank sum | 0.021 (After ACV+light - Baseline)<br>0.0079 (After light - After ACV+light) |
| **Fig. 4i** | Trial#<br>PAM12>CsC: | Baseline moving trials | kruskal wallis | 4.7E-5 (PAM12>CsC)<br>0.306 (UAS-CsC) |
|  | Baseline: 12<br>After ACV+light: 55<br>After light: 50<br>from 22 flies<br>UAS-CsC:<br>Baseline: 8<br>After ACV+light: 42<br>After light: 40<br>from 21 flies |  |  |  |
|  |  |  | Post hoc<br>bonferroni | PAM12>CsC:<br>5.31E-5 (Baseline vs After ACV+light)<br>0.0455 (After ACV+light vs After light)<br>0.0164 (Baseline vs After light)<br>UAS-CsC:<br>0.725 (Baseline vs After ACV+light)<br>1 (After ACV+light vs After light)<br>0.38 (Baseline vs After light) |
| **Fig. 4f**<br>**(top)** | Trial# 30<br>from 12 flies | Flies without<br>alerting, non-alerting and<br>spontaneous<br>locomotion;<br>Baseline<br>moving trials | Wilcoxon<br>signed<br>rank | 0.56 |
| **Fig. 4f**<br>**(bottom)** | Trial# 31<br>from 16 flies | Flies without<br>alerting, non-alerting and<br>spontaneous<br>locomotion;<br>Baseline<br>moving trials | Wilcoxon<br>signed<br>rank | 0.65 |
| **Fig. 4h**<br>**(top)** | Fly#<br>PAM12>GtA: 11<br>UAS-GtA: 17 | None | Wilcoxon<br>rank sum | 0.013 |
| **Fig. 4h**<br>**(bottom);**<br>**Extended**<br>**Data Fig.**<br>**7g** | Fly#<br>PAM12>GtA: 17<br>PAM12-GAL4: 13 | None | Wilcoxon<br>rank sum | Trial 1: 0.0941<br>Trial 2: 0.357<br>Trial 3: 0.0065<br>Trial 4: 0.0074<br>Trial 5: 0.005 |
| **Extended**<br>**Data Fig.**<br>**7b** | Fly#<br>PAM12>CsC: 7<br>PAM13, 14>CsC: 7<br>UAS-CsC: 9 | None | Wilcoxon<br>signed<br>rank | PAM12>CsC:<br>0.0156 (Baseline vs After ACV+light)<br>0.0156 (After ACV+light vs After light)<br>PAM13, 14>CsC:<br>0.0156 (Baseline vs after ACV+light)<br>0.688 (After ACV+light vs After light)<br>UAS-CsC:<br>0.0039 (Baseline vs after ACV+light)<br>0.652 (After ACV+light vs After light) |
| **Extended**<br>**Data Fig.**<br>**7c** | Fly#<br>PAM12>CsC: 7<br>PAM13, 14>CsC: 7<br>UAS-CsC: 9 | None | Wilcoxon<br>rank sum | PAM12>CsC vs UAS-CsC:<br>0.031 (After ACV+light - Baseline)<br>0.0021 (After light - After ACV+light)<br>PAM13, 14>CsC vs UAS-CsC:<br>0.68 (After ACV+light - Baseline)<br>0.68 (After light - After ACV+light) |
| **Extended Data Fig. 7e** | Fly#<br>PAM12>CsC: 15<br>PAM12-GAL4: 15<br>UAS-CsC: 16 | None | kruskal wallis | 0.663 (Before PAM12 activation)<br>0.0075 (After PAM12 activation) |
|  |  |  | post hoc bonferroni | Before PAM12 activation:<br>1 (PAM12>CsC vs PAM12-GAL4)<br>1 (PAM12>CsC vs UAS-CsC)<br>1 (PAM12-GAL4 vs UAS-CsC)<br>After PAM12 activation:<br>0.176 (PAM12>CsC vs PAM12-GAL4)<br>0.00569 (PAM12>CsC vs UAS-CsC)<br>0.708 (PAM12-GAL4 vs UAS-CsC) |
| **Extended Data Fig. 7f (top left)** | Fly#<br>PAM12>GtA: 11<br>UAS-GtA: 17 | None | Wilcoxon rank sum | 0.0055 |
| **Extended Data Fig. 7f (top right)** | Fly#<br>PAM12>GtA: 11<br>UAS-GtA: 17 | None | Wilcoxon rank sum | ACV + light<br>Trial 1: 1<br>Trial 2: 0.888<br>Trial 3: 0.0428<br>Trial 4: 0.0144<br>Trial 5: 0.0127<br>Trial 6: 0.0014<br>Trial 7: 0.0022<br>Trial 8: 0.0041<br>Trial 9: 0.0035<br>Trial 10: 0.11 |
| **Extended Data Fig. 7f (bottom left)** | Fly#<br>PAM13, 14>GtA: 7<br>UAS-GtA: 10 | None | Wilcoxon rank sum | 0.54 |
| **Extended Data Fig. 7f (bottom center)** | Fly#<br>PAM13, 14>GtA: 7<br>UAS-GtA: 10 | None | Wilcoxon rank sum | ACV + light<br>Trial 1: 0.669<br>Trial 2: 0.813<br>Trial 3: 0.193<br>Trial 4: 0.109<br>Trial 5: 0.962<br>Trial 6: 0.74<br>Trial 7: 0.364<br>Trial 8: 0.887<br>Trial 9: 0.315<br>Trial 10: 0.536 |
| **Extended Data Fig. 7f (bottom right)** | Fly#<br>PAM13, 14>GtA: 7<br>UAS-GtA: 10 | None | Wilcoxon rank sum | 0.67 |
| **Fig. 4j** | Trial#<br>MBON09>CsC:319<br>MBON09-GAL4:292<br>UAS-CsC:363<br>Fly#<br>MBON09>CsC:60<br>MBON09-GAL4:54 | Baseline non-grooming trials | Fisher's exact test | 1.15E-24 (MBON09>CsC vs MBON09-GAL4)<br>2.35E-15 (MBON09>CsC vs UAS-CsC)<br>0.0045 (MBON09-GAL4 vs UAS-CsC) |
|  | UAS-CsC:65 |  |  |  |
| **Fig. 4k** | Fly#<br>PAM12>GtA: 39<br>PAM12-GAL4: 39<br>UAS-GtA: 38 | Baseline non-grooming trials | Wilcoxon rank sum | 0.00233 (PAM12>GtA vs PAM12-GAL4)<br>0.00775 (PAM12>GtA vs UAS-GtA)<br>0.984 (UAS-GtA vs PAM12-GAL4) |
| **Fig. 4l** | Trial#<br>Alerting: 92<br>Non-alerting: 183<br>from 31 flies | Baseline moving trials | Wilcoxon rank sum | 1.39E-7 (Preceding moving distance)<br>0.348 (Preceding time) |
| **Fig. 5d** | Fly#<br>MBON09>CsC: 12<br>MBON09-lexA: 9 | None | Wilcoxon rank sum | 0.036 |
| **Fig. 5h** | Trials#<br>Offset response +:<br>47<br>-: 69<br>from 21 flies | Baseline moving trials, non-alerting trials | Wilcoxon rank sum | 0.004 |
| **Fig. 5k (light)** | Fly#<br>PAM>CsC: 10<br>PAM-lexA: 10 | None | Wilcoxon rank sum | 0.026 |
| **Fig. 5k (offset)** | Fly#<br>PAM>CsC: 10<br>MBON09>CsC: 12 | None | Wilcoxon rank sum | 0.0011 |
| **Fig. 5j** | Trial# (10 trials/fly)<br>MBON21>GtA: 216<br>MBON21-GAL4: 183<br>UAS-GtA: 172 | Baseline non-grooming trials; Non-alerting trials | Kruskal-Wallis | 6.72E-12 |
|  |  |  | Post hoc Bonferroni | 1.2E-7 (MBON21>GtA vs MBON21-GAL4)<br>2.32E-10 (MBON21>GtA vs UAS-GtA)<br>1 (MBON21-GAL4 vs UAS-GtA) |
| **Fig. 5i** | Trial#<br>MBON21>CsC: 65<br>MBON21-GAL4: 72<br>UAS-CsC: 65<br>Fly#<br>MBON21>CsC: 22<br>MBON21-GAL4: 17<br>UAS-CsC: 20 | Non-alerting trials, baseline moving trials | Wilcoxon rank sum | 0.0231 (UAS-CsC vs MBON21-GAL4)<br>3.66E-5 (MBON21>CsC vs MBON21-GAL4)<br>0.043 (MBON21>CsC vs UAS-CsC) |
| **Extended Data Fig. 9d** | Fly#<br>MBON09>CsC: 24<br>UAS-CsC: 24<br>MBON09-GAL4: 24 | Baseline moving trials | Wilcoxon signed rank | Average speed over 5s: 0.0086 (MBON09>CsC)<br>0.75 (MBON09-GAL4)<br>0.55 (UAS-CsC)<br>Locomotor duration:<br>0.0079 (MBON09>CsC)<br>0.64 (MBON09-GAL4)<br>0.67 (UAS-CsC) |
| **Extended Data Fig. 9e** | Fly #<br>MBON21>GtA: 22<br>UAS-GtA: 22<br>MBON21-GAL4: 20 | None | Kruskal-Wallis | 0.452 |
|  |  |  | Post hoc<br>Bonferroni | 0.64 (MBON21>GtA vs MBON21-GAL4)<br>1 (MBON21>GtA vs UAS-GtA) |
| **Extended**<br>**Data Fig.**<br>**9f** | Fly #<br>MBON21>CsC: 21<br>MBON21-GAL4:<br>20<br>UAS-CsC: 19 | Baseline<br>moving trials | Wilcoxon<br>rank sum | 0.0354 (MBON21>CsC vs MBON21-GAL4)<br>0.0219 (MBON21>CsC vs UAS-CsC)<br>0.622 (MBON21-GAL4 vs UAS-CsC) |
| **Extended**<br>**Data Fig.**<br>**9h** | Trial#<br>MBON21>GtA: 216<br>MBON21-GAL4:<br>183<br>UAS-GtA: 172<br>Fly#<br>MBON21>GtA: 26<br>MBON21-GAL4:<br>19<br>UAS-GtA: 20 | Baseline<br>non-grooming<br>trials | Fisher's<br>exact test | 0.0793 (MBON21-GAL4 vs UAS-GtA)<br>0.0435 (MBON21-GAL4 vs MBON21>GtA)<br>0.000167 (UAS-GtA vs MBON21>GtA) |
| **Fig. 6f** | Trials#<br>baseline response<br>+: 63<br>-: 76<br>from 37 flies | Baseline<br>moving<br>trials, non-alerting trials | Wilcoxon<br>rank sum | 0.0089 |
| **Fig. 6i** | Fly #<br>CRE011>Kir: 9<br>UAS-Kir: 8 | None | Wilcoxon<br>rank sum | Mean peak amplitude: 0.015<br>Peak No. per min: 0.0023<br>Integral: 0.0055 |

